# The phase separation-dependent FUS interactome reveals nuclear and cytoplasmic function of liquid-liquid phase separation

**DOI:** 10.1101/806158

**Authors:** Stefan Reber, Helen Lindsay, Anny Devoy, Daniel Jutzi, Jonas Mechtersheimer, Michal Domanski, Oliver Mühlemann, Silvia M.L. Barabino, Marc-David Ruepp

## Abstract

Liquid-liquid phase separation (LLPS) of proteins and RNAs has emerged as the driving force underlying the formation of membrane-less organelles. Such biomolecular condensates have various biological functions and have been linked to disease. One of the best studied proteins undergoing LLPS is Fused in Sarcoma (FUS), a predominantly nuclear RNA-binding protein. Mutations in *FUS* have been causally linked to Amyotrophic Lateral Sclerosis (ALS), an adult-onset motor neuron disease, and LLPS followed by aggregation of cytoplasmic FUS has been proposed to be a crucial disease mechanism. In spite of this, it is currently unclear how LLPS impacts the behaviour of FUS in cells, e.g. its interactome. In order to study the consequences of LLPS on FUS and its interaction partners, we developed a method that allows for the purification of phase separated FUS-containing droplets from cell lysates. We observe substantial alterations in the interactome of FUS, depending on its biophysical state. While non-phase separated FUS interacts mainly with its well-known interaction partners involved in pre-mRNA processing, phase-separated FUS predominantly binds to proteins involved in chromatin remodelling and DNA damage repair. Interestingly, factors with function in mitochondria are strongly enriched with phase-separated FUS, providing a potential explanation for early changes in mitochondrial gene expression observed in mouse models of ALS-FUS. In summary, we present a methodology that allows to investigate the interactome of phase-separating proteins and provide evidence that LLPS strongly shapes the FUS interactome with important implications for function and disease.

## Introduction

The biophysical process of liquid-liquid phase separation (LLPS) has drawn considerable attention over the last couple of years. Indeed, the regulators as well as the biophysical driving forces of LLPS are only just beginning to be understood. LLPS was not only reported to be important for the formation of various membraneless organelles such as nucleoli, P-bodies and stress granules, but also has been implicated in various diseases (Alberti and Carra, 2018; Alberti and Dormann, 2019; Boeynaems et al., 2018). One of the best studied proteins that undergoes LLPS *in vitro* and *in vivo* is Fused in Sarcoma (FUS) (Hofweber et al., 2018; Kang et al., 2019; Kato et al., 2012; Monahan et al., 2017; Murray et al., 2017; Patel et al., 2015; Schwartz et al., 2013). FUS is an ubiquitously expressed RNA-binding protein that has been implicated in diverse RNA metabolic pathways, such as transcription, pre-mRNA splicing and miRNA processing (Meissner et al., 2003; Raczynska et al., 2015; Reber et al., 2016; Schwartz et al., 2012; Yu and Reed, 2015; Zhang et al., 2018). In 2009, mutations in *FUS* were shown to be causative for Amyotrophic Lateral Sclerosis (ALS) (Kwiatkowski et al., 2009; Vance et al., 2009). ALS is the most common motor neuron disease in human adults and is characterised by a progressive loss of upper and lower motor neurons, causing paralysis and ultimately leading to death (Cleveland and Rothstein, 2001). Most ALS mutations in FUS disrupt its C-terminal nuclear localisation signal (NLS), leading to cytoplasmic mislocalisation and aggregation of FUS in neurons and glial cells of affected individuals, a pathological hallmark of FUS-ALS (Dormann et al., 2010; Ling et al., 2013). Several recently published mouse models indicate a toxic gain of FUS function in the cytoplasm (Devoy et al., 2017; Scekic-Zahirovic et al., 2016; Sharma et al., 2016). Noteworthy, recruitment of FUS into phase-separated RNP granules, e.g. stress granules, followed by aggregation has been proposed to drive disease (Alberti and Hyman Anthony, 2016; Wolozin and Ivanov, 2019).

Due to the current lack of tools to study membraneless organelles (Tang, 2019), LLPS-dependent FUS interactions are unknown and it is unclear how cytoplasmic FUS exerts its toxic function(s). Furthermore, it is also unknown if and how phase separation contributes to FUS function. Aiming to better understand functional consequences of FUS phase separation and to identify FUS interactors under LLPS conditions, we developed a method that allows for the purification of phase-separated FUS together with its associated proteins and RNAs and compared these interactors to FUS interactors that were purified under non-LLPS conditions. We observed distinct interaction patterns depending on the biophysical state of FUS. Whereas FUS binds predominantly to factors involved in RNA processing under non-LLPS conditions, phase separated FUS preferably binds to proteins involved in chromatin remodeling, DNA damage response and proteins with functions in mitochondria. Furthermore, our data suggest that LLPS in different cellular compartments has different functions. While phase separation is required for binding of FUS to chromatin and for at least some of its functions in the nucleus, LLPS in the cytoplasm appears not to be required for FUS toxicity. Possibly, phase separation followed by aggregation might be the pathway through which cells cope with increased cytoplasmic FUS in order to reduce the concentration of soluble FUS and its deleterious function(s). In summary, our method provides new insights into how LLPS affects FUS interaction partners and function. Notably, this method should be applicable to other proteins that undergo LLPS and allows to address the importance of LLPS on their function and interactome.

## Results

### Determining the liquid-liquid phase separation dependent FUS interactome

In order to determine which protein and RNA species interact with FUS under LLPS conditions, we developed a novel approach which allows for the purification of phase separated FUS from cell lysates (summarised in Fig. 1a). Enhanced green fluorescent protein (eGFP)-tagged wild type FUS or FUS harbouring the ALS-associated mutation P525L, which disrupts the function of the C-terminal NLS (Dormann et al., 2010), were transiently expressed in HEK293T cells (Fig. 1b). After cell lysis, the volume of the lysate was reduced to approximately half of its original volume. This volume reduction resulted in the formation of eGFP-FUS-containing droplets within the cell lysate, which can be visualised by fluorescence microscopy (Supplementary Fig. 1a). To stabilise these FUS-droplets, the sample was treated with the reversible crosslinker formaldehyde. Thereafter, FUS-droplets were stable enough to be analysed by flow cytometry (Supplementary Fig. 1b) and to be sorted by fluorescence activated particle sorting (Supplementary Fig. 1c). To address which protein and RNA species interact with FUS under non-LLPS conditions, a regular co-immunoprecipitation (co-IP) was performed using nanobodies against eGFP. As phase separation of FUS is highly dependent on FUS concentration (Patel et al., 2015; Wang et al., 2018) and the wash volumes applied during the co-IP exceeded the volume applied to analyse the cell lysates in Supplementary Fig. 1b, LLPS of FUS during co-IP conditions is limited to the minimum or even completely prevented. A pulldown of FLAG-eGFP served as control IP. Successful purification of the bait from droplet purifications and co-IPs were verified by SDS-PAGE followed by silver staining or western blotting, respectively (Supplementary Fig. 2). Proteins and RNAs purified from co-IP and droplet purification experiments were analysed by means of quantitative mass spectrometry or RNA deep sequencing, respectively. Interestingly, the wild type and P525L FUS protein and RNA interactomes (under the same experimental conditions) were mostly identical (Supplementary Fig. 3), which is why wild type and P525L interactomes from the same experimental conditions were pooled, in order to identify the most robust FUS interactors under LLPS and non-LLPS conditions. We identified 238 proteins interacting with FUS under LLPS conditions and 360 under non-LLPS conditions. 102 proteins were present in both datasets (independent of either biophysical state), resulting in 136 proteins specific to the LLPS condition. The observation that several proteins and RNAs preferentially interacted with FUS under LLPS conditions (Fig. 1c and Supplementary Table 1) indicates that altered biophysical conditions within phase separated droplets enable FUS to undergo novel interactions. Of note, while many proteins did not pass the statistical criteria (see material and methods) to be assigned to both interactomes, many proteins assigned to either the LLPS or the non-LLPS interactome were detected in the reciprocal experiment. Indeed, 70 % of the LLPS interactors were detected in the non-LLPS condition, while 81 % of the non-LLPS interactors were detected in the LLPS experiment. This indicates different affinities, but not necessarily mutually exclusive binding to FUS, depending on its biophysical state. We further analysed proteins co-purified with FUS under LLPS and non-LLPS conditions, comparing them to previously reported FUS interactors (Blokhuis et al., 2016; Chi et al., 2018; Hein et al., 2015; Kamelgarn et al., 2016; Reber et al., 2016; Sun et al., 2015; Wang et al., 2015). Interestingly, only 23.8 % of the LLPS-dependent FUS interactors have been previously reported, whereas 47.2 % of the co-IPed FUS interactors have already been reported by these previous studies. This is in line with the idea that phase separation changes the affinity of FUS for its interaction partners. As aforementioned studies used non-LLPS conditions (co-immunoprecipitations) to purify FUS and its interaction partners, it is not surprising that there are clearly more previously unknown FUS interactors present in the LLPS-dependent FUS interactome.

**Figure 1.**
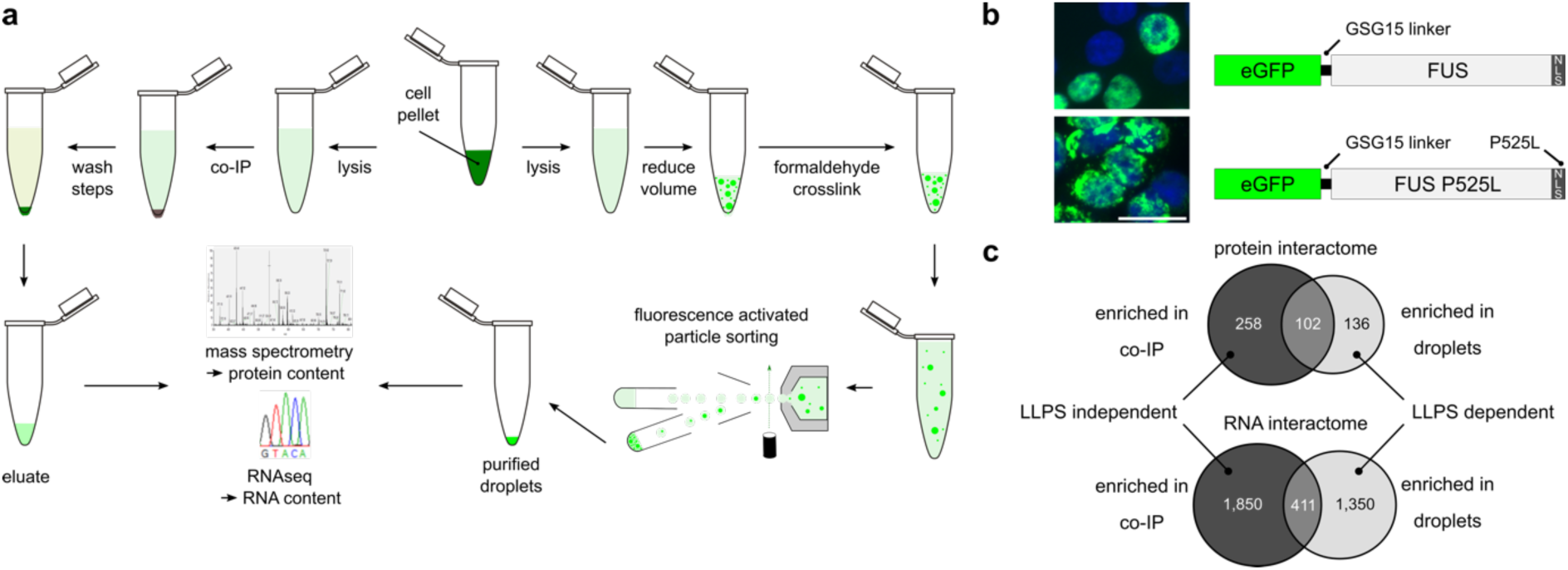
Co-IP and purification of LLPS FUS followed by quantitative mass spectrometry and RNA deep sequencing. **a** Experimental workflow. HEK293T cells expressing wild type or P525L eGFP-FUS fusion protein are lysed and subsequently subjected either to a co-IP experiment using anti-GFP nanobodies coupled to magnetic beads (left path) or to eGFP-FUS droplet purification (right path). Droplets are generated through reducing the volume of the lysate and stabilized using the reversible crosslinker formaldehyde. Thereafter, the droplets are purified by fluorescence activated particle sorting and additional wash steps. **b** Constructs used for co-IP and droplet purification experiments. eGFP fused to FUS including a GSG15 linker between the two proteins. Wild type eGFP-FUS (top right) localizes mainly to the nucleus whereas ALS mutant P525L eGFP-FUS (bottom right) localizes predominantly to the cytoplasm as shown by fluorescence microscopy of transiently transfected HeLa cells counterstained with DAPI. Scale bar = 30 µm. **c** Summary of quantitative mass spectrometry (top) and RNA deep sequencing (bottom) experiments. Shown are numbers for protein and RNA species which were significantly enriched in co-immunopurification and droplet purification experiments comprising the respective overlap between the two datasets.

While, compared to the number of significantly enriched protein interactors, higher numbers of RNAs could be identified in FUS droplets (1,761) or co-IPed together with FUS (2,261), with 411 RNAs significantly enriched in both samples (Fig. 1c and Supplementary Table 2), the overall picture of the RNA interactome is similar to the protein interactome. More interactors were identified in the co-IP compared to the droplet purification experiment, and the relative overlap between the two samples is similar for both, protein and RNA interactomes.

### FUS has a different protein interactome depending on its biophysical state

In order to identify protein families enriched in LLPS and non-LLPS conditions, we performed a protein network analysis using STRING (Szklarczyk et al., 2015) on the 136 proteins that were exclusively co-purified with FUS under LLPS conditions and compared them to all the proteins (n = 360) that were co-IPed together with FUS (non-LLPS conditions). Interestingly, STRING classified LLPS-specific FUS interactors into three main functional groups (Fig. 2a), namely into proteins with functions in mitochondria (red), proteins involved in chromatin remodelling and DNA damage response (blue) and proteins involved in RNA splicing (green). Although enriched in the LLPS-specific FUS interactome, proteins involved in RNA splicing were much more prominent in the STRING generated for proteins co-immunoprecipitated together with FUS (Fig. 2b, green). In line with this finding, performing gene ontology (GO) term enrichment analyses on the same groups of proteins using the WEB-based GEne SeT AnaLysis Toolkit (WebGestalt) (Liao et al., 2019) revealed that factors involved in RNA splicing and mRNA processing were enriched much more significantly in the non-LLPS FUS interactome compared to the LLPS-specific FUS interactome (Supplementary Fig. 4a).

**Figure 2.**
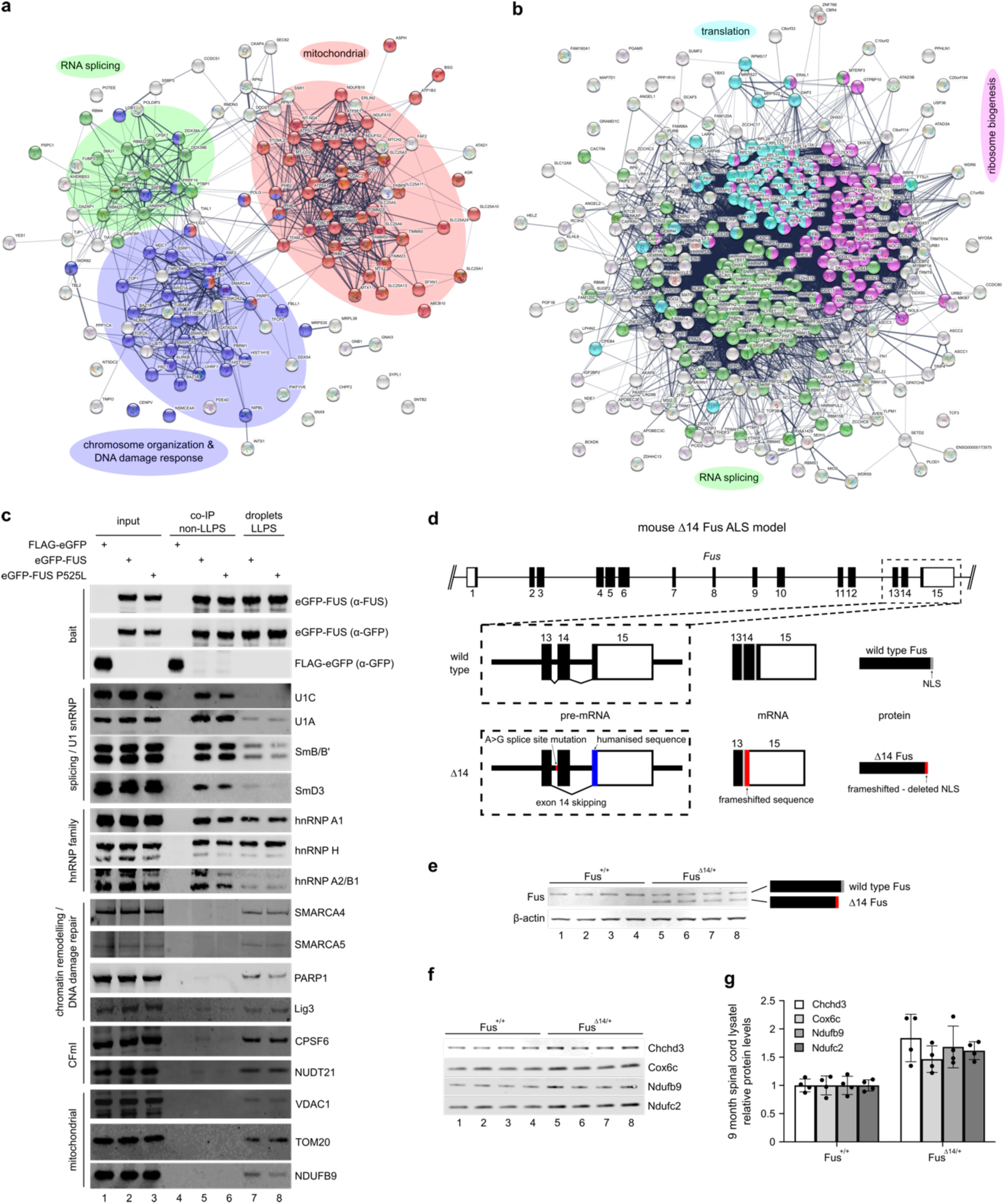
Preferential protein interaction partners depending on FUSs biophysical state and dysregulated mitochondrial protein homeostasis at an early stage in FUS-ALS. **a** STRING analysis of LLPS-specific FUS interactors (n=136). Highlighted are proteins with biological functions in RNA splicing (green, GO:0008380), chromosome organization and cellular response to DNA damage stimulus (blue, GO:0051276 and GO:0006974) and proteins mitochondrial functions (red, GO:0006839, GO:0007005 and GO:0006811). **b** STRING analysis of non-LLPS interactors of FUS (n=360). Highlighted are proteins with biological functions in RNA splicing (green, GO:0008380), translation (cyan, GO:0006412) and ribosome biogenesis (magenta, GO:0042254). **c** Western blot analysis of proteins co-immunoprecipitated (non-LLPS) with control (FLAG-eGFP, lane 4) or FUS (lane 5-6) and purified together with FUS droplets (LLPS, lane 7-8). **d** Scheme of the ‘FUSDelta14’ knockin mouse ALS model. A reported ALS mutation (FUS p.G466VfsX14) destroys the 3’ splice site of exon 14 leading to exon skipping resulting in a novel C-terminus deleting the endogenous FUS NLS. To generate the identical frameshift peptide to that of the human patient, human exon 15 coding sequence was also knocked-in. **e** Western blot analysis of spinal cord lysates from 9 month old FUS^+/+^ (lane 1-4) and FUSΔ^14/+^ (lane 5-8) mice. While FUS^+/+^ mice only express full-length Fus, FUSΔ^14/+^ mice express full-length and the shorter Δ14 Fus. β-actin served as loading control. **f** As in e, but showing changes in mitochondrial protein levels. Protein levels were normalized to total protein levels in each sample. **g** Quantification of western blot in f showing protein levels relative to wild type normalized to total protein levels.

To validate the proteins detected by mass spectrometry, we performed western blot analysis of proteins involved in RNA splicing, chromatin remodelling and DNA damage repair and factors with functions in mitochondria (Fig. 2c). While some proteins showed the same binding to FUS regardless of LLPS, such as hnRNP H and hnRNP A1, others showed a clear preference for either LLPS or non-LLPS conditions. Consistent with the mass spec data (Supplementary Table 1), proteins involved in mRNA splicing were co-purified with FUS independent of its biophysical state. Nonetheless, they seemed to interact with FUS preferentially under non-LLPS conditions. Furthermore, proteins involved in chromatin remodelling and DNA damage response as well as mitochondrial proteins were almost exclusively detectable together with phase-separated FUS. Interestingly, nuclear FUS granules have already been reported to associate with RNA polymerase II (RNA Pol II) and the N-terminus of FUS, which can phase-separate and is also the transcriptional activator domain of FUS-CHOP and FUS-ERG fusion proteins observed in cancer, is sufficient to target the SWI/SNF chromatin remodelling complex (Burke et al., 2015; Kato et al., 2012; Linden et al., 2019; Prasad et al., 1994; Thompson et al., 2018; Zinszner et al., 1994). Moreover, chromatin remodelling is an important aspect of DNA damage response and FUS granules have already been reported at sites of DNA damage (Patel et al., 2015). Therefore, it makes sense that proteins involved in DNA damage response were specifically enriched with phase separated FUS. Strikingly, we also detected the members of cleavage factor I (CFIm) in our LLPS FUS interactome. Besides its function in mRNA 3’-end processing, CFIm has been linked to chromatin remodelling (Yu et al., 2018).

The most unexpected and at the same time prominent LLPS-dependent FUS interactors were proteins with function in mitochondria. Although cytoplasmic FUS was previously reported to interact with mitochondria (Deng et al., 2015), it was surprising that mitochondrial proteins formed the top GO term in the LLPS-specific FUS interactome (Supplementary Fig. 4a). Intriguingly, it has previously been reported that dysregulation in mitochondrial gene expression occurs at the initial disease stage in ‘FUSDelta14’ knockin mice, which heterozygously express FUS carrying a C-terminal frameshift mutation and hence deleted NLS (Devoy et al., 2017). To assess if mitochondrial protein levels are affected at an early, pre-symptomatic stage in the ‘FUSDelta14’ mouse model (Fig. 2d and e), we quantified proteins isolated from spinal cord sections of pre-symptomatic mice by western blotting (Fig. 2f and g) normalizing to total protein levels in each sample (Supplementary Fig. 4b). While proteins involved in RNA processing, DNA damage repair and chromatin remodelling appeared unchanged (Supplementary Fig. 4c), several mitochondrial proteins were dysregulated in the ‘FUSDelta14’ knockin mice. Of note, this was not due a general change of mitochondrial homeostasis, as other mitochondrial proteins remained unaffected (Supplementary Fig. 4d). These data suggest that phase separation increases the affinity of FUS towards mitochondria in the cytoplasm, potentially increasing the local FUS concentration around mitochondria, resulting in deleterious effects.

### FUS has a different RNA interactome depending on its biophysical state

While mRNAs were the most abundant RNAs associated with either FUS droplets (LLPS conditions) or non-phase separated FUS (co-IP) (Fig. 3a-c), the most prominently enriched RNA species co-purified with FUS under both conditions were U snRNAs (Fig. 3d and e). U snRNAs are the RNA components of small nuclear ribonuclear particles (snRNPs) and are responsible for recognition and removal of introns from pre-mRNAs during splicing (Jutzi et al., 2018). Although this enrichment could be expected for RNAs co-purified with non-phase separated FUS, as proteins involved in pre-mRNA splicing are also strongly enriched under non-LLPS conditions, the strong enrichment of snRNAs together with phase-separated FUS was rather surprising, as snRNP protein components are not significantly enriched together with LLPS FUS. Interestingly, cytoplasmic FUS granules have been previously reported to mislocalize U snRNAs (Gerbino et al., 2013; Reber et al., 2016). Indeed, we could detect U1 and U11 snRNAs together with ALS-linked FUS-P525L in the cytoplasm, whereas U1 snRNP specific proteins remained nuclear (Fig. 3g). Together, this data suggests that phase separated FUS binds preferentially to naked snRNAs, while non-phase separated FUS binds mainly to fully assembled snRNPs and functions in splicing. Indeed, we and others have already shown that FUS contacts both the major (U2- type) and minor (U12-type) spliceosome to regulate splicing of specific introns (Reber et al., 2016; Sun et al., 2015; Yu and Reed, 2015). Finally, the biggest difference between the two purification experiments was the strong enrichment of mitochondrial ribosomal RNAs (MtrRNAs) (Fig. 2f), which were strongly deriched in the non-LLPS condition, but clearly enriched in the LLPS condition. Importantly, this is consistent with the proteomic data, where we detected high levels of mitochondrial proteins purified together with FUS droplets, but not in the co-IP condition.

**Figure 3.**
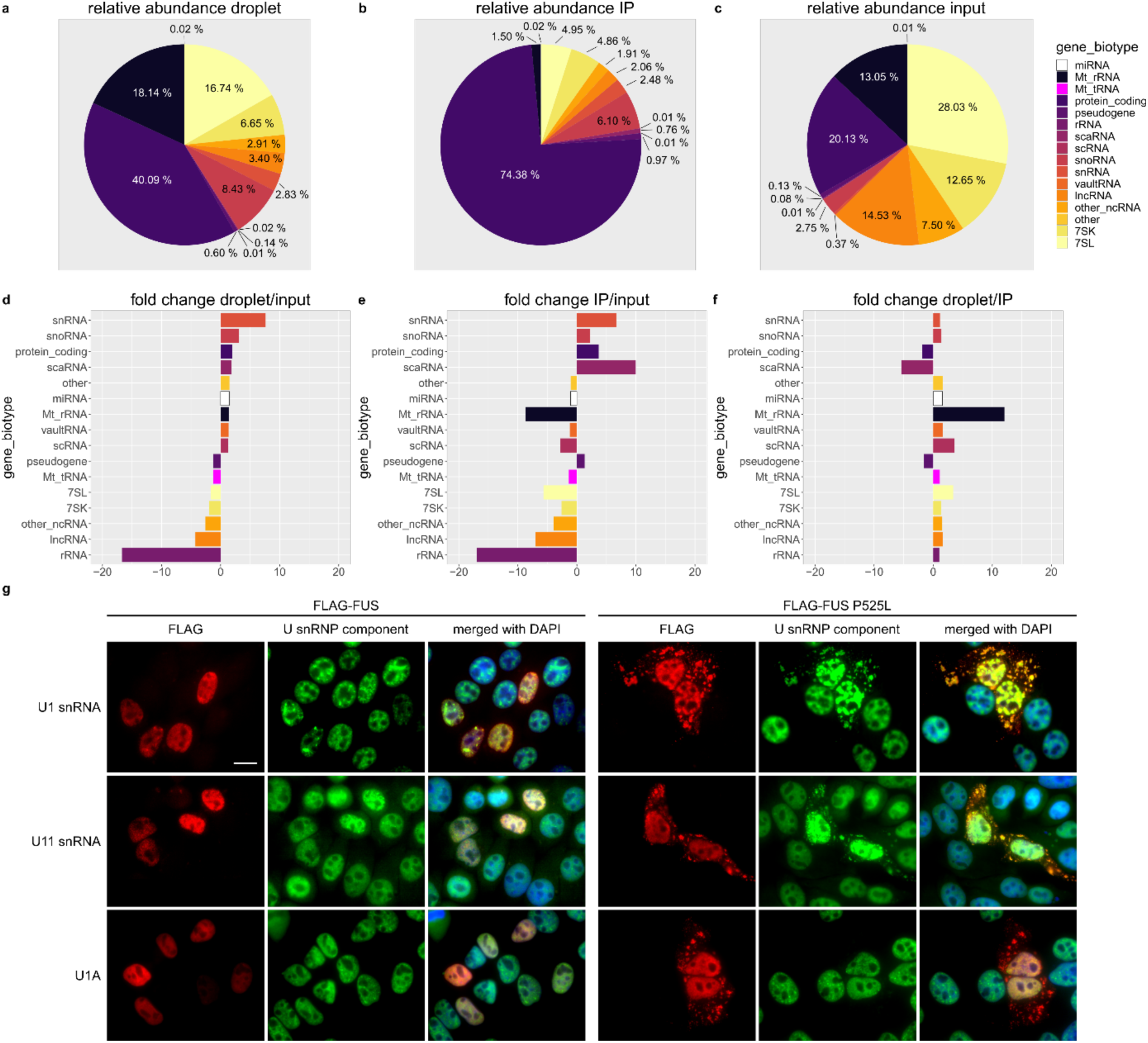
RNA interactome of LLPS and non-LLPS FUS. **a** Relative abundance of different RNA species co-purified with LLPS FUS (FUS droplets). **b** Relative abundance of different RNA species co-purified with non-LLPS FUS (co-IP). **c** Relative abundance of different RNA species in the input sample. **d** fold change of relative RNA abundance of different RNA species comparing RNAs found under LLPS conditions relative to the input. **e** Same as in d, but comparing RNA found in non-LLPS conditions compared to the input. **f** Same as in d, but comparing RNA abundance between LLPS and non-LLPS conditions. **g** HeLa cells transiently transfected with either FLAG-FUS (left) or FLAG-FUS P525L (right) and stained for FLAG (red channel) and different components of the U1 snRNP (green channel), either by RNA-FISH (U1 and U11, first and second row) or immunostaining (U1A, third row). Cells were counterstained using DAPI. Scale bar = 15 µm.

### LLPS is required for the association of FUS with chromatin and its function in autoregulation

In order to assess the importance of FUS LLPS for FUS function, we created an N-terminally FLAG-tagged FUS construct and substituted 27 tyrosines in the N-terminal prion-like domain (PLD) with serines (PLD27YS FUS). The aromatic ring structures of the tyrosines in the FUS PLD were previously shown to drive LLPS through inter-and intermolecular cation-π interactions with positively charged amino acid side chains (Qamar et al., 2018; Wang et al., 2018). Mutating these tyrosines to serines abolishes phase separation *in vitro* and *in vivo* (Kato et al., 2012; Wang et al., 2018). Strikingly, PLD27YS FUS showed slightly increased cytoplasmic localization compared to wild type FUS (Fig. 4a, first two rows). This suggests that besides the C-terminal NLS of FUS, also the N-terminus contributes to nuclear localization of FUS at steady state, possibly through phase-separation-dependent nuclear interactions of FUS. Indeed, it was previously reported that the N-terminus of FUS is required for binding of FUS to chromatin (Yang et al., 2014), where FUS localises to granules and regulates gene expression in a transcription-dependent manner. Moreover, inhibition of transcription leads to cytoplasmic re-localisation of FUS, suggesting that active transcription tethers FUS to newly synthesized RNA bound to chromatin (Patel et al., 2015; Thompson et al., 2018; Zinszner et al., 1994). This is consistent with the recent finding that FUS leaves the nucleus through passive diffusion (Ederle et al., 2018). If LLPS is indeed required for FUS binding to chromatin, one would expect more FUS diffusing from the nucleus to the cytoplasm if phase separation is inhibited. To be able to study the importance of LLPS for FUS function in the nucleus and to exclude that a loss of function is not due to FUS mislocalisation, we generated a phase separation-deficient FUS construct with an additional strong SV40 NLS to ensure nuclear localization (Fig 4a, last row). To investigate the importance of LLPS for the ability of FUS to bind to chromatin, we transiently expressed wild type and LLPS-deficient FUS in HEK293T cells, followed by a cytoplasmic/nucleoplasmic vs chromatin biochemical fractionation. Indeed, LLPS-deficient FUS hardly interacted with chromatin compared to the wild type FUS (Fig. 4b and c), indicating that phase separation is required for FUS function in co-transcriptional gene expression.

**Figure 4.**
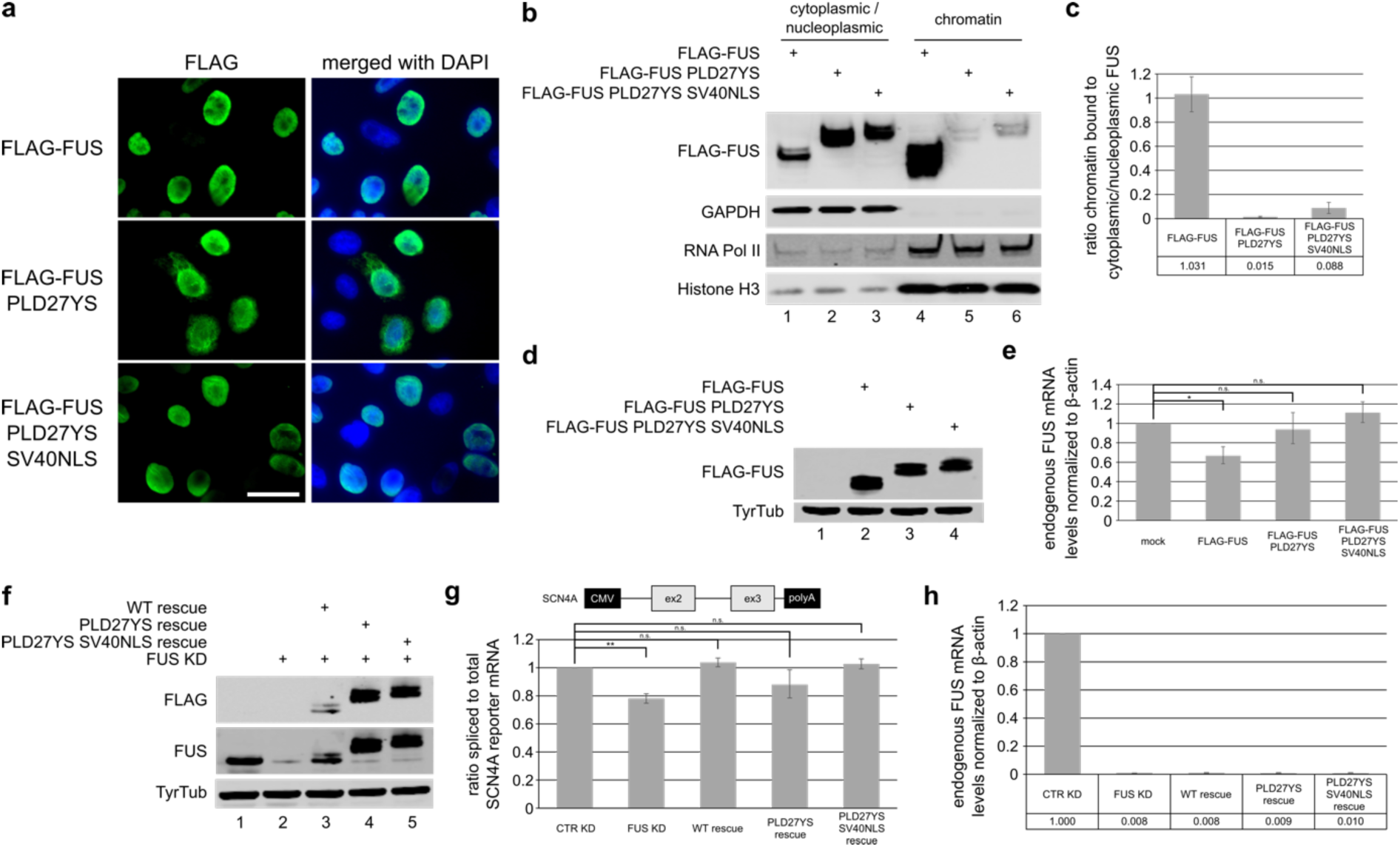
Phase separation is required for FUS binding to chromatin and function in FUS autoregulation. **a** Immunostaining of HeLa cells transiently expressing wild type FLAG-FUS, or LLPS deficient FLAG-FUS PLD27YS and FLAG-FUS PLD27YS SV40NLS, respectively. Scale bar = 30 µm. **b** Western blot of cytoplasmic/nucleoplasmic vs chromatin fractionation experiment of HEK293T cells transiently expressing the constructs used in a. The blots were incubated with antibodies against FLAG. GAPDH, RNA Pol II and Histone H3 serve as controls for the respective fractions. While wild type FLAG-FUS is strongly bound to chromatin together, phase separation deficient FUS is almost absent in the chromatin fraction. **c** Quantification of western blots in b. Shown is the ratio of chromatin bound to cytoplasmic/nucleoplasmic FUS relative to the wild type FLAG-FUS construct. Average values and standard deviation of three biological replicates are shown. **d** Western blot of HeLa cells which were either mock, FLAG-FUS, FLAG-FUS PLD27YS or FLAG-FUS PLD27YS SV40NLS transfected. Total cell lysates were subjected to SDS-PAGE and western blotting with antibodies against FLAG. Tyrosine tubulin served as loading control. **e** Endogenous FUS mRNA levels as determined by RT-qPCR relative to mock transfected cells. Average and standard deviations of three biological replicates are shown. Single asterisk indicates a p-values of < 0.05. **f** Western blot analysis of FUS levels under control knockdown (CTR KD, lane 1), FUS KD (lane 2) and FUS rescue (lanes 3-5) conditions using wild type and LLPS deficient FUS constructs. Proteins from HeLa extracts were separated by SDS-PAGE and blots were incubated with anti-FLAG (upper row) and anti-FUS (middle row). Tyrosine tubulin (lower row) serves as loading control. **g** Ratio of spliced to total RNA expressed from the SCN4A minigene (The minigene is driven by a CMV promoter and expresses exon 2 and 3 and the intervening U12-type intron) under CTR KD, FUS KD and FUS KD followed by a rescue with different RNAi-resistant expression constructs. Average values and standard deviations of three biological replicates are shown. Double asterisk indicates a p-values of < 0.01. **h** Relative endogenous FUS mRNA levels from samples analysed in g.

To test if LLPS-deficient FUS is still functional, we made use of two previously established assays for FUS function. First, we tested if PLD27YS FUS is still capable of autoregulating endogenous FUS mRNA levels when transiently transfected into HeLa cells (Loughlin et al., 2019). While wild type FUS autoregulated endogenous FUS mRNA levels, LLPS-deficient FUS had no effect on endogenous FUS mRNA levels (Fig 4d and e). Importantly, this was not due cytoplasmic mislocalisation of PLD27YS FUS as the addition of the SV40 NLS did not rescue FUS function in autoregulation (Fig. 4d and e). Next, we used the minor (also referred to as U12-type) intron containing SCN4A minigene, which requires FUS for efficient splicing (Reber et al., 2016). If cells are depleted of FUS, splicing of the SCN4A minigene becomes less efficient and this effect can be rescued by transient expression of RNAi-resistant FUS. Surprisingly, LLPS-deficient FUS was still able to promote efficient splicing (Fig 4f and g), even though it was unable to autoregulate. While PLD27YS FUS was only partially active, PLD27YS FUS harbouring the additional SV40 NLS fully rescued splicing of the SCN4A minigene. This difference presumably occurs due to the partial cytoplasmic mislocalisation of PLD27YS FUS, which is rescued upon the addition of an additional NLS. Importantly, these differences did not emerge from differences in knockdown efficiencies of endogenous FUS, which are identical between the experimental conditions (Fig. 4h). The capability of LLPS-deficient FUS to promote splicing is in line with our observation that non-phase-separated FUS preferentially interacts with proteins involved in RNA splicing, indicating that phase separation is not necessary for FUS function in splicing.

### LLPS is not required for FUS toxicity in the cytoplasm

Proteins with functions in mitochondria were highly enriched in our LLPS-dependent FUS interactome. Besides our observation that mitochondrial proteins were already dysregulated in pre-symptomatic ALS-FUS mice, others reported that cytoplasmic FUS has deleterious effects on mitochondria leading to apoptosis (Cha et al., 2019; Deng et al., 2015; Stoica et al., 2016). Therefore, we examined the impact of LLPS on FUS-mediated toxicity via mitochondria. We transiently expressed FLAG-tagged wild type FUS, P525L FUS and LLPS-deficient P525L FUS in HEK293T cells using the previously introduced PLD27YS mutant. To assess FUS toxicity, we measured the release of cytochrome c from mitochondria to the cytoplasm, a key step during the mitochondrial apoptotic pathway (Wang and Youle, 2009), by performing mitochondrial versus cytoplasmic biochemical fractionations. To exclude potential toxic effects of the transfection reagent used for this experiment, untreated cells were compared to mock transfected cells, excluding a toxic effect of the transfection reagent (Supplementary Fig. 5a and b). As expected, overexpression of either wild type or P525L FUS caused toxicity (Fig. 5a and b), in line with previous studies reporting that overexpression of FUS has deleterious effects in different models (Lindstrom and Liu, 2018; Mitchell et al., 2013; Sabatelli et al., 2013; Steyaert et al., 2018). Interestingly, LLPS-deficient cytosolic FUS also exhibited a toxic effect, suggesting that phase separation is not required for FUS toxicity.

**Figure 5.**
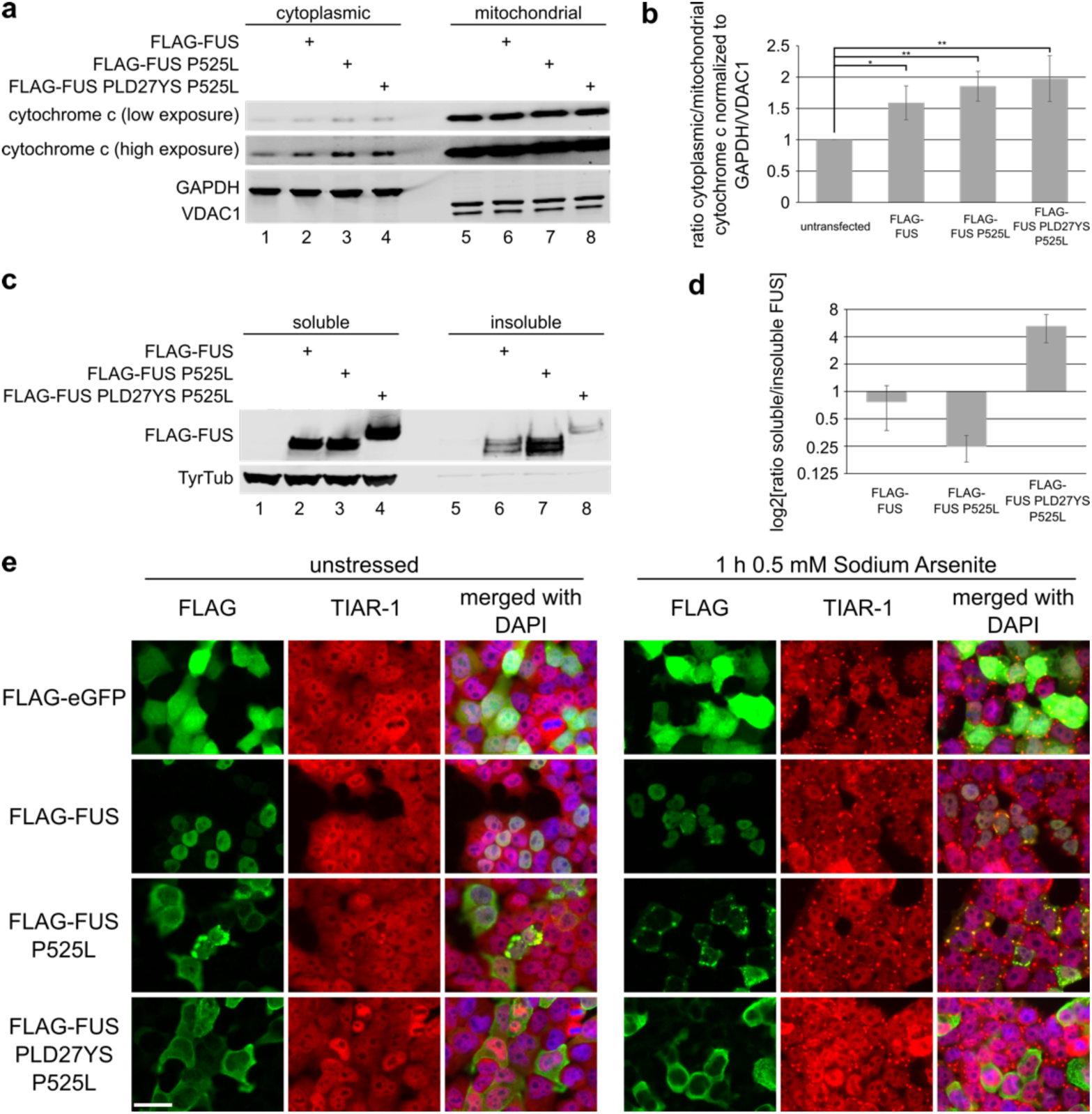
LLPS is not required for cytoplasmic FUS toxicity. **a** Cytoplasmic/mitochondrial fractionation of untransfected, FLAG-FUS, FLAG-FUS P525L and FLAG-FUS PLD27YS P525L transfected HEK293T cells, respectively. Cytoplasmic (lanes 1-4) and mitochondrial (lanes 5-8) fractions were analysed by western blotting using anti cytochrome c antibody (top and middle row). Cytochrome c signal is shown in two different exposures. GAPDH and VDAC1 (lower row) served as controls for cytoplasmic and mitochondrial fractions, respectively. **b** Quantification of cytochrome c levels in a. Shown are the ratios of cytoplasmic to mitochondrial cytochrome c relative to the control. Average values and standard deviations from five biological replicates are shown. Single and double asterisk indicate a p-values of < 0.05 and < 0.01, respectively. **c** Western blot of soluble/insoluble fractionation of the cells transfected in a. Membranes were incubated with anti-FLAG antibodies. Tyrosine tubulin served as a loading control. **d** Quantification of FUS levels in c. Shown is the ratio of soluble to insoluble FUS relative to wild type FLAG-FUS. Average values and standard deviation from five biological replicates are shown. **e** Immunostaining of unstressed (left) and stressed (right) HEK293T cells transiently transfected with the indicated FLAG-FUS constructs. In addition, a FLAG-eGFP control was included. Cells were stained for FLAG (green) and the stress granule marker TIAR-1 (red). Cells were counterstained using DAPI. Scale bar = 30 µm.

As FUS aggregates were suggested to exert a toxic function, we compared the solubility of LLPS-competent and LLPS-deficient FUS. To this end, half of the cells transfected for the mitochondrial/cytoplasmic fractionation were subjected to a RIPA soluble/insoluble biochemical fractionation experiment. As previously reported (Coady and Manley, 2015), ALS mutant cytoplasmic FUS exhibited reduced solubility compared to wild type FUS (Fig. 5c and d). Interestingly, LLPS-deficient cytoplasmic FUS was more soluble than both wild type and cytoplasmic LLPS-competent FUS. Indeed, it was already shown, using purified FUS protein *in vitro,* that phase separation precedes the formation of insoluble aggregates (Patel et al., 2015). Our data strongly suggests that also in more complex environments, such as cell systems and *in vivo,* phase separation is involved in the formation of insoluble FUS aggregates, and therefore may be the mechanism by which FUS aggregates observed in post-mortem tissue of ALS-FUS patients are formed.

It has been postulated that recruitment of FUS to stress granules (SGs) may be the driving force in the processes from cytoplasmic mislocalisation to insoluble aggregation (Dormann et al., 2010; Ling et al., 2013). Indeed, SGs are phase separated cytoplasmic compartments (Hyman et al., 2014), and various studies reported FUS localization to SGs (Bosco et al., 2010; Dormann et al., 2010; Hock et al., 2018; Matus et al., 2014). In order to investigate the effect of LLPS on SG recruitment of FUS, we transiently transfected HEK393T cells with the constructs described above and induced oxidative stress using sodium arsenite, which was reported to efficiently recruit cytoplasmic, but not nuclear FUS, to SGs (Bentmann et al., 2012; Bosco et al., 2010; Dormann et al., 2010; Hock et al., 2018). As an additional control, FLAG-eGFP was transfected. As expected, cytoplasmic P525L FUS robustly localized to arsenite induced SGs together with the stress granule marker TIAR-1 (Fig. 5e). Notably, even under unstressed conditions, P525L FUS containing stress granules were detectable in some cells, presumably due to stress induced through P525L FUS overexpression or reduced chaperoning of FUS-P525L by TNPO1 (Hofweber et al., 2018) and hence enhanced SG recruitment. Furthermore, some wild type FUS expressing cells showed co-localization of FUS together with stress granule markers under stress, likely due to the very high FUS levels leading to cytoplasmic FUS in these cells. Strikingly, cells expressing LLPS-deficient FLAG-FUS PLD27YS P525L did not (or to a very low extent) form stress granules under oxidative stress, in contrast to neighbouring non-transfected cells that form stress granules normally (Fig. 5e).

It is known that stress granule formation does not require FUS (Aulas et al., 2012). To exclude the possibility that LLPS-deficient FUS only prevents TIAR-1 from being recruited to stress granules, but not stress granule formation per se, we performed the same experiment with immunostaining for G3BP1, another well-defined stress granule marker (Aulas et al., 2012). As observed for TIAR-1, also G3BP1 wasn’t recruited to SGs during oxidative stress if LLPS-deficient FUS was present, whereas it co-localized to stress granules with LLPS-competent P525L FUS (Supplementary Fig. 5c). To further exclude that this behaviour is specific to arsenite-induced oxidative stress, we also applied osmotic stress to HEK293T cells transiently expressing the aforementioned FUS constructs using D-sorbitol. Consistent with the oxidative stress condition, LLPS-deficient FUS also prevented the formation of stress granules under osmotic stress (Supplementary Fig. 5d). There are two possible explanations for this behaviour: Either LLPS-deficient FUS protects cells from oxidative and osmotic stress, or LLPS-deficient FUS prevents stress granule formation through sequestration of factors required for this process. Although the first possibility cannot be excluded at this point it seems rather unlikely, especially since LLPS-deficient FUS is at least as toxic as LLPS-competent FUS in our experiments. Interestingly, we (this study, see Fig. 2 (PARP1) and Supplementary Table 1) and others (Blokhuis et al., 2016; Bosco et al., 2010; Kamelgarn et al., 2016; Sun et al., 2015) reported interactions between FUS and different stress granule markers. Hence, it is conceivable that LLPS-deficient FUS sequesters factors required for the formation of stress granules and thus prevents the formation of stress granules during stress. In summary, we provide evidence that LLPS and aggregation of FUS are not required for FUS toxicity on mitochondria. Nonetheless, LLPS seems to be important for the recruitment of FUS to stress granules and potentially proceeding to formation of insoluble FUS.

## Discussion

In this study, we describe a new method that allows for the purification of liquid-liquid phase separated proteins combining chemical crosslinking with fluorescence activated particle sorting. This approach allowed us to identify new and validate previously reported FUS interactors. Furthermore, we show that LLPS changes the FUS interactome, presumably due to different affinities of FUS to its interactors depending on its biophysical state. Nevertheless, many FUS interactors are detected under both conditions but show a clear preference for either dispersed/soluble or phase separated FUS.. Importantly, several proteins enriched with phase separated FUS, such as DDX3X, DHX9, FMR1, TIAR- 1 and SMN1 (compare Supplementary Table 1), have already been observed by others to co-localize with FUS into cytoplasmic granules (Blokhuis et al., 2016; Bosco et al., 2010; Groen et al., 2013; Kamelgarn et al., 2016). Moreover, proteins found in nuclear granules, specifically paraspeckles (Shelkovnikova et al., 2014) and transcription-dependent granules containing FUS and RNA Pol II (Thompson et al., 2018) are highly enriched under LLPS conditions compared to non-LLPS conditions (compare data in Supplementary Table 1).

We identified factors involved in chromatin remodelling and DNA damage repair to be the most prominent nuclear protein families binding with high preference to LLPS FUS. Interestingly, numerous recent studies reported that transcription factors recruit the mediator coactivator complex through phase separation using their activation domains leading to recruitment of RNA Pol II binding RNA Pol II C-terminal domain (CTD) (Boehning et al., 2018; Boija et al., 2018; Cho et al., 2018; Chong et al., 2018; Sabari et al., 2018). Noteworthy, FUS was previously reported to interact with the CTD of RNA Pol II (Schwartz et al., 2012) and was identified in transcription-dependent granules together with RNA Pol II (Thompson et al., 2018). Moreover, the N-terminal domain of FUS, which was identified as the transcriptional activator in FUS-CHOP and FUS-ERG fusion proteins in liposarcoma and myeloid sarcoma respectively (Prasad et al., 1994; Zinszner et al., 1994), was recently shown to be sufficient to contact the SWI/SNF chromatin remodelling complex (Linden et al., 2019). Strikingly, protein components of the mediator complex as well as protein subunits of RNAP II detected by mass spectrometry are clearly more abundant under LLPS conditions. Indeed, many of these proteins were exclusively detected in the LLPS condition while they were completely absent under non-LLPS conditions (Supplementary Fig. 6). In addition, components of the mammalian pre-mRNA 3’ end processing factor CFIm (composed of NUDT21 (also CPSF5), CPSF6 and CPSF7) which were linked to chromatin remodelling (Yu et al., 2018), are much stronger enriched together with LLPS FUS compared to non-LLPS FUS, while in contrast, the other members of the 3’-end processing machinery are slightly higher enriched under non-LLPS conditions (Supplementary Fig. 7). In line with the idea that phase separation of FUS plays an important role in FUS chromatin-associated function(s), we show that FUS requires LLPS to efficiently bind to chromatin. Consistently, LLPS deficient FUS mostly dissociates from chromatin and partially re-localizes to the cytoplasm. Most likely, LLPS FUS binds to chromatin in a transcription-dependent manner, since previous studies reported re-localization of FUS to the cytoplasm upon transcription inhibition (Patel et al., 2015; Zinszner et al., 1994) and FUS was found in transcription-dependent granules together with RNA Pol II (Thompson et al., 2018). Our data provides further evidence that LLPS is the driving force for FUS binding to chromatin and the cytoplasmic re-localisation of LLPS deficient FUS, as a consequence of losing its nuclear tether, is consistent with recent data (Ederle et al., 2018), which suggests that FUS leaves the nucleus predominantly through passive diffusion In addition, we show that LLPS of FUS is required for FUS autoregulation, indicating that during transcription FUS is recruited through LLPS to the *FUS* gene to regulate its own expression. Nonetheless, LLPS is not required to promote splicing of the minor intron containing SCN4A reporter gene. On the one hand, this is might be explained by our observation that FUS binds to factors involved in splicing preferentially under non-LLPS conditions. On the other hand, the intron of the SCN4A minigene is most likely spliced after transcription termination. Indeed, minor introns are typically spliced slower post-transcriptionally compared to their major counterparts (Patel et al., 2002; Singh and Padgett, 2009; Younis et al., 2013). Consequently, FUS and the minor spliceosome probably bind to the SCN4A pre-mRNA after transcription termination in a non-LLPS state. Of note, it has recently been shown that TDP-43, another protein of the hnRNP family that undergoes LLPS, does to require phase separation to perform its function in pre-mRNA splicing (Schmidt et al., 2019). This is in line with studies which performed *in vitro* splicing assays in dependence on FUS or TDP-43, respectively (Deshaies et al., 2018; Meissner et al., 2003). Both studies used HeLa nuclear extracts where nuclear components are highly diluted (compared to the *in vivo* context) and thus phase separation of either FUS or TDP-43 very unlikely. Nonetheless, both proteins function in splicing in the *in vitro* context. Altogether, these findings strongly indicate that LLPS is not required for the function of neither FUS nor TDP-43 in pre-mRNA splicing.

Besides providing evidence for the importance of FUS LLPS for FUS nuclear function in autoregulation, our data suggests that LLPS and aggregation of FUS are not required for FUS toxicity on mitochondria. Of note, recent ALS-FUS mouse models, including the FUSDelta14 mouse that express cytoplasmic Fus from the endogenous mouse locus or at endogenous levels, consistently showed motor neuron degeneration in the absence of FUS aggregation. Indeed, all of these studies failed to detect cytoplasmic FUS inclusion bodies which are observed in human tissue (Devoy et al., 2017; Scekic-Zahirovic et al., 2016; Sharma et al., 2016). Although toxicity of cytoplasmic FUS aggregates cannot be excluded at this point, these findings strongly indicate that increased cytoplasmic concentrations of FUS are sufficient for having deleterious effects, leading to motor neuron death. Moreover, our data suggests that FUS LLPS is not necessary for the toxic effects of cytoplasmic FUS. It is therefore tempting to speculate that the formation of insoluble cytoplasmic FUS inclusions could be a mechanism how cells get rid of increasing amounts of cytoplasmic FUS. Namely through local concentration of FUS, leading to FUS LLPS and subsequent recruitment to SGs, followed by precipitation through a liquid-to-solid state transition. Thereby, cells might prevent (or reduce) deleterious effects, reducing the amount of soluble (toxic) FUS in the cytoplasm. Of note, this is not a new concept in the field of neurodegeneration: similarly, amyloid plaques in Alzheimer’s disease have been proposed to buffer toxic soluble amyloid-beta species (Brody et al., 2017). While it remains to be seen if LLPS and aggregation are dispensable for cytoplasmic FUS toxicity *in vivo*, our data provides evidence that LLPS is indeed the precursor of insoluble FUS aggregates, as LLPS deficient FUS is clearly more soluble than LLPS competent FUS when overexpressed in HEK293T cells. This indicates that similar to *in vitro* studies (Patel et al., 2015), also *in vivo* LLPS is a prerequisite for liquid-to-solid state transition of FUS leading to the formation of insoluble FUS aggregates. Additionally, LLPS deficient FUS does not localize to oxidative or osmotic stress induced stress granules and even, at least partially, inhibits stress granule formation, while LLPS competent FUS robustly localizes to stress granules.

To conclude, we present a method which allows to identify LLPS-specific protein and RNA interactors of a protein of interest which should be applicable to other proteins undergoing LLPS. We find that LLPS alters and expands the protein and RNA interactome of FUS. In the nucleus, LLPS is important for anchoring FUS to the chromatin and proper nuclear localization at steady state and favoring interactions with the chromatin remodeling and DNA damage repair machineries. Furthermore, our data suggest that LLPS in the cytoplasm is not a prerequisite for FUS to exert its toxicity.

## Acknowledgements

This research and related results were made possible through the support of the NOMIS Foundation (MDR), the support by the UK Dementia Research Institute which receives its funding from DRI Ltd, funded by the UK Medical Research Council, Alzheimer’s Society and Alzheimer’s Research UK (MDR), the Motor Neurone Disease Association (AD; 867-791), and the support of the NCCR RNA and Disease funded by the Swiss National Science Foundation (MDR, OM). We also thank, Muriel Fragnière, Sabrina Schenk and Tosso Leeb of the Next Generation Sequencing platform (University of Bern) for technical advice and library preparation. Moreover, we thank Sophie Marie-Pierre Braga, Anne-Christine Uldry and Manfred Heller of the Proteomics Mass Spectrometry Core Facility (University of Bern) for technical advice and sample processing. We further thank Dorothee Dormann for critical reading of the manuscript, and Nicole Kleinschmidt, Karin Schranz, and Joana Conde de Almeida de Sousa Furtado for their excellent technical and organisational support.

## Data availability

Mass spectrometry and high-throughput sequencing data will be accessible upon paper acceptance.

## Authors Contributions

S.R. and M.-D.R. designed and conceived the study. S.R. performed experiments and analysis shown in figures 1 supported by M.D., 2a-c, 3a-f, 4 and 5 and the associated supplementary figures. H.L. performed the computational analysis of the RNA-Seq data. A.D. performed experiments shown in figure 2d-g and S4b-d. D.J. and J.M. performed FISH and IF experiments in figure 3g and S5d. S.R. wrote the manuscript with inputs from M.-D.R. A.D., O.M., S.M.L.B. and M.-D.R. critically revised the manuscript. All authors contributed to the discussion and data interpretation and the final version of the manuscript.

## Methods

### Oligonucleotides, Plasmids, Antibodies

Oligonucleotides, plasmids and antibodies are described in the Supplementary Information.

### Cell culture

HeLa and HEK293T cells were maintained in Dulbecco’s modified Eagle’s medium (DMEM) containing 10 % fetal calf serum (FCS), penicillin (100 U/µl) and streptomycin (100 µg/ml)) at 37 °C and 5 % CO_2_. Cells were transfected using Dogtor (OZ Biosciences) for all experiments except for cytoplasmic/mitochondrial (membrane), soluble/insoluble fractionation experiments and transfections of HEK293T cells prior to stress followed by immunofluorescence. For these experiments, cells were transfected using TransIT-LT1 (Mirus).

### Generation of cell lysates for droplet purification and eGFP co-immunoprecipitation

30 µg plasmid coding for eGFP-GSG15-FUS or 15 µg plasmid coding for FLAG-eGFP, respectively were transiently transfected into 40-50 % confluent HEK293T cells in a T300 flask using Dogtor according to the manufacturer’s instructions. The medium was replaced 24 h after transfection. 48 h after transfection, the cells were harvested by flushing with ice cold medium. Cells were pelleted at 4 °C at 200 g for 5 min. After removal of the supernatant, the cells were washed with ice cold PBS. Thereafter, cells were pelleted at 4 °C for 3 min at 1’500 g. PBS was removed and the cells were snap frozen in liquid nitrogen and stored at −80 °C until usage. The pellet was thawed on ice and re-suspended in 400 µl lysis buffer (75 mM HEPES pH7.3, 100 mM KOAc, 0.5 mM DTT, 0.5 % NP40, 1:5’000 Antifoam B, 10 µl/ml lysis buffer protease inhibitor, 10 µl/ml lysis buffer RNase inhibitor) and transferred to a 1.5 ml Eppendorf tube. To increase lysis efficiency, the cells were passed 5 x through a 25G 5/8 needle. The lysate was subsequently used to isolate FUS droplets or to perform an eGFP co-immunoprecipitation.

### Purification of FUS-containing droplets

To increase the formation of LLPS FUS droplets, the volume of the lysate was reduced to approximately half of the initial volume in a speedvac. To reduce time in the speedvac, samples were split into several smaller samples of approx. 150 µl. To monitor droplet formation, a drop of the lysate was placed on a standard microscopy glass slide, covered with a coverslip and FUS droplets were visualized using a fluorescence microscope. Subsequently, 30 µl concentrated lysate were transferred to a new 1.5 ml Eppendorf tube and droplets were stabilized by adding 0.3 µl 10 % formaldehyde (final conc. 0.1 %). The lysate was mixed by vortexing and incubated for 8 min at room temperature. Remaining cross-linker was quenched through the addition of 1 µl 1 M TRIS pH7.3 and vortexing. The sample was thereafter stored on ice until droplets were sorted by fluorescence activated particle sorting (FAPS). Right before sorting, 300 µl PBS was added to the lysate and the sample was passed through a 40 µm cell strainer (to get rid of aggregates which could clog the sorter). Droplets were sorted into PBS according to eGFP-fluorescence and side scatter (SSC) on a FACS ARIA (BD Biosciences). Sorted droplets were stored at – 80 °C until further processing. Before protein or RNA isolation, the droplets were pelleted by centrifugation at 4 °C at 16’000 g for 15 min. The droplet-pellet was washed with 1 ml PBS. The wash was repeated one more time. Between washes, the droplets were centrifuged for 15 min 4 °C at 16’000 g. After removal of the PBS, 150’000 droplets (sorted events) were either re-suspended in 50 µl 1X LDS-loading buffer (NuPAGE™ LDS Sample Buffer (4X) (NP0007, Thermo Fisher) diluted to 1X supplemented with 50 mM DTT) for protein elution or in 50 µl RNA-sample buffer (50 mM TRIS pH7.0, 5 mM EDTA, 1 % SDS, 10 mM DTT) for subsequent RNA isolation.

### eGFP Nanobodies and coupling to magnetic beads

Plasmids encoding for His-tagged anti-GFP nanobodies (clones LaG-9, LaG-16 and LaG-24) were obtained under the MTA from Michael P Rout laboratory (Rockefeller University, New York, USA). All three constructs were expressed in ArticExpress (DE3) cells (Agilent) and purified using Ni-NTA resin as described (Fridy et al., 2014). Purified anti-GFP nanobodies were coupled to magnetic beads (Dynabeads M-270 Epoxy, Invitrogen) accordingly to manufacturer’s instruction. The coupling mixture contained 20 μg of purified nanobody per 1 mg of beads. The coupling reaction was carried out (separately for each clone) with rotation, at 37°C for 20 hours. Anti-GFP nanobodies coupled to magnetic beads were re-suspended in 50% glycerol/PBS (2 mL per 300 mg beads) and stored at − 20°C. In order to increase GFP binding efficiency, beads coupled to 3 different clones were mixed together in 1:1:1 ratio.

### eGFP co-immunoprecipitation

20 µl GFP nanobodies coupled to magnetic beads were transferred to 1 ml lysis buffer (75 mM HEPES pH7.3, 100 mM KOAc, 0.5 mM DTT, 0.5 % NP40, 1:5’000 Antifoam B, 10 µl/ml lysis buffer protease inhibitor, 10 µl/ml lysis buffer RNase inhibitor). Lysis buffer was removed and washed again with another ml lysis buffer. After removal of the lysis buffer, the beads were re-suspended in 100 µl lysis buffer and added to 400 µl previously prepared cell lysate. The beads were incubated for 3 h at 4 °C head over tail on a rotor. Thereafter, the supernatant was removed and the beads were washed 5 x 5 min in 1 ml of 2 x lysis buffer at 4 °C head over tail on a rotor. The first wash step contained protease and RNase inhibitors. After the last wash step, the beads were splitted in two parts and either re-suspended in 1 ml TRIZOL for RNA isolation or 50 µl 1X LDS-loading buffer for protein elution.

### Mass spectrometry

The formaldehyde crosslink was reversed / eGFP-fusion proteins were eluted from the beads by boiling the samples for 15 min at 95 °C in LDS-loading buffer. Cell lysates mixed with equal volumes of 2X LDS-loading buffer and boiled for 15 min at 95 °C served as input samples. To prepare the samples (each experimental condition in biological triplicates) for mass spectrometry, samples were run 1 cm into a 12 % Bis-Tris Plus gel (Invitrogen). 150’000 droplets (sorted events) and 3/5 of the eGFP co-IP, respectively were loaded. The gel was subsequently washed 5 x 5 min in ulatapure water, 10 min in 0.1 M HCl and Coomassie stained for 2 h (0.12 % (w/v) Coomassie G-250, 10 % H_3_PO_4_ (v/v), 10 % (w/v) NH_4_OAc, 20 % MeOH (v/v)) to visualize proteins. The gel was de-stained for 8 x 30 min in ultrapure water. To prepare gel pieces for mass spectrometry, the gel was cut, using a clean scalpel, into approx. 0.25 cm horizontal bands. The bands were further cut into cubes of approx. 1-3 mm^3^ and transferred to a 1.5 ml Eppendorf tube. Samples were stored at 4 °C in 20 % EtOH before reduction and alkylation. Samples were analyzed in a random order to avoid chromatographic batch effects. The gel pieces were reduced, alkylated and digested by trypsin as described elsewhere (Gunasekera et al., 2012). The digests were analyzed by nano-liquid chromatography tandem masspectrometry (nLC-MS/MS) (EasyLC 1000 nanoflow-UPLC coupled to a QExactive HF mass spectrometer, ThermoFisher Scientific) with one injection of 5 μl digests. Peptides were trapped on a Precolumn (C18 PepMap100, 5 μm, 100 Å, 300 μm × 5 mm, Thermo Fisher Scientific, Reinach, Switzerland) and separated by backflush on a C18 column (3 μm, 100 Å, 75 μm × 15 cm, C18, Nikkyo Technos, Tokyo, Japan) by applying a 40-minute gradient of 5 % acetonitrile to 40 % in water, 0.1 % formic acid, at a flow rate of 350 nl/min. The Full Scan method was set with resolution at 60,000 with an automatic gain control (AGC) target of 1E06 and maximum ion injection time of 50 ms. The data-dependent method for precursor ion fragmentation was applied with the following settings: resolution 15,000, AGC of 1E05, maximum ion time of 110 milliseconds, isolation width of 1.6 m/z, collision energy 27, under fill ratio 1 %, charge exclusion of unassigned and 1+ ions, and peptide match preferred, respectively. Spectra interpretation was performed with MaxQuant/Andromeda version 1.5.4.1 searching against the forward and reversed SwissProt Homo Sapiens protein database (Release 2017_12) using fixed modification of carbamidomethylation on Cys, and variable modifications of oxidation on Met, deamidation on Asn/Gln, and acetylation on protein N-term. Mass error tolerance for parent ions was set to 10 ppm, fragment ion tolerance to 20 ppm, and full trypsin cleavage specificity with 3 missed cleavages were allowed. Based on reversed database matches a 1 % false discovery rate (FDR) was set for acceptance of peptide spectrum matches (PSM), peptides, and proteins. Relative protein abundance was calculated as described elsewhere (Krey et al., 2014). In brief, contaminant proteins such as keratins and trypsin were removed and the remaining protein’s iBAQ values were each divided by the sum of all non-contaminant iBAQ values. Relative iBAQ values were used to determine fold changes and p-values using student’s t-test and adjusted p-values (FDR) between experimental conditions. To be assigned to the LLPS interactome of FUS, proteins purified from FUS droplets had to be enriched > 2-fold compared to the input with a FDR < 0.05. To be assigned to the non-LLPS interactome, proteins co-IPed with FUS had to be enriched > 2-fold compared to the input as well as to the control IP (FLAG-eGFP), with a FDR < 0.05 for both.

### RNA isolation and RNA deep sequencing

To isolate RNA from the droplets, the formaldehyde crosslink was reversed through incubation of the droplets in RNA-sample buffer (50 mM TRIS pH7.0, 5 mM EDTA, 1 % SDS, 10 mM DTT) for 40 min at 70 °C. After cooling of the sample on ice for 5 min, 1 ml TRIZOL was added. RNA was isolated from TRIZOL according to the manufacturer’s instructions. Quality and quality of RNA was analyzed with an Agilent 2100 Bioanalyzer (Agilent Technologies). Total RNA isolated from transiently transfected cells served as input RNA. Total RNA isolated from droplets, from the co-IP and input RNA (in biological triplicates for each sample) were ribodepleted using the RiboMinus™ Transcriptome Isolation Kit (Invitrogen, K155004) before library preparation according to the manufacturer’s manual. Libraries were prepared using the strand-specific Illumina TruSeq Stranded Total RNA kit (Part # 15031048 Rev. E). Total RNA libraries were sequenced on the Illumina HiSeQ3000 platform using 100 bp single-end sequencing cycles. Adapters and low quality bases were trimmed from reads using TrimGalore v0.4.4 (Krueger, 2015; Martin, 2011). Reads were then mapped using Salmon v0.8.2 (Patro et al., 2017) to the human cDNA and non-coding RNA transcriptome, ENSEMBL version 38.90. Transcripts per million (TPM) were imported into R using tximport v1.4.0 (Soneson et al., 2015), and differential gene expression analysis performed using edgeR v3.18.1 (Robinson et al., 2010). Transcript lengths were included as an offset in modelling, and transcript length scaled TPMs were used for calculating average log2(TPM). RNAs with > 2-fold change to the input with a FDR < 0.001 were considered as significantly enriched over the input. To assess relative abundances of different RNA biotypes, several ENSEMBL gene biotypes were summarized in groups (pseudogenes include: *“transcribed_unitary_pseudogene”, “unprocessed_pseudogene”, “processed_pseudogene”, “transcribed_unprocessed_pseudogene”, “polymorphic_pseudogene”, “transcribed_processed_pseudogene”, “IG_V_pseudogene”, “unitary_pseudogene”, “TR_V_pseudogene”, “TR_J_pseudogene”, “IG_C_pseudogene”, “IG_J_pseudogene”, “translated_processed_pseudogene”, “pseudogene”, “IG_pseudogene“*. <u>lncRNAs include</u>: *“antisense_RNA”, “lincRNA”, “sense_intronic”, “sense_overlapping”, “bidirectional_promoter_lncRNA”, “3prime_overlapping_ncRNA”, “macro_lncRNA“*. other_ncRNA include: *“processed_transcript”, “misc_RNA”, “ribozyme”, “sRNA”, “non_coding“*. other include: *“TR_V_gene”, “IG_V_gene”, “IG_C_gene”, “IG_J_gene”, “TR_J_gene”, “TR_C_gene”, “IG_D_gene” “TR_D_gene”*. 7SL and 7SK genes were excluded from the above groups to form each a separate group.

### STRING and Gene ontology (GO) enrichment analysis

STRING (Szklarczyk et al., 2015) analysis was performed using the multi protein search function using default options. The “confidence” option was chosen to indicate strength of data support for each interaction. In addition, GO terms indicated in the figure legends were highlighted. GO term analysis was performed using the WEB-based GEne SeT AnaLysis Toolkit (WebGestalt) (Liao et al., 2019) using default settings with the following options: Organism of Interest: Homo sapiens; Method of Interest: Over-Representation Analysis (ORA); Functional Database: geneontology, Biological Process noRedundant. For GO term analysis of genes identified by mass spectrometry, the Reference Set “genome protein-coding” was used. For GO term analysis of genes identified by RNAseq, the Reference Set “genome” was used.

### Flow cytometry to analyze formation of FUS-droplets

To analyze FUS droplets by flow cytometry, FUS droplets were prepared as described above. Droplets were analyzed on a LSR II SORP H274 [BD Biosciences] and generated data was processed using FlowJo version 10.

### Cytoplasmic vs mitochondrial (membrane) and soluble vs insoluble fractionations

60-80 % confluent HEK293T cells in T25 flasks were harvested using Trypsin/EDTA and washed once with ice cold PBS. The cells were split in two parts. One part was lysed in RIPA buffer (Thermo Fisher, 89900) containing 1X Halt™ Protease Inhibitor Cocktail (Thermo Fisher, 78429) for 20 min on ice with occasional vortexing. Samples were centrifuged for 15 min at 16’000 g at 4 °C and the supernatant was subsequently mixed with equal volumes of 2X LDS-loading buffer (NuPAGE™ LDS Sample Buffer (4X) (NP0007, Thermo Fisher) diluted to 2X supplemented with 100 mM DTT) and boiled for 5 min at 95 °C (soluble fraction). The insoluble pellet was washed twice with PBS and thereafter incubated in isoluble buffer (50 mM Tris pH7.5, 200 mM NaCl, 2 mM KCl, 1 mM EDTA, 0.5 % Glycerol, 100 mM Urea, 1X Halt™ Protease Inhibitor Cocktail) for 1 h at 37 °C at 1,400 rpm. The sample was centrifuged for 5 min at 16,000 g and the supernatant was subsequently mixed with equal volumes of 2X LDS-loading buffer and boiled for 5 min at 95 °C (insoluble fraction). The other part of the cells was used for cytoplasmic/mitochondrial fractionation according to (Baghirova et al., 2015) with slight modifications: Cells were re-suspended in 400 µl buffer A (150 mM NaCl, HEPES pH7.5, 25 µg/ml Digitonin (Sigma Aldrich, D141), 1 M Hexylene glycol (Sigma Aldrich, 112100), 1X Halt™ Protease Inhibitor Cocktail) and incubated for 10 min at 4 °C head over tail on a rotor. Samples were centrifuged for 10 min at 2,000 g at 4 °C and the supernatant (cytoplasmic fraction) was transferred to a new 2 ml Eppendorf tube. The pellet was subsequently washed 2 x with buffer A (5 min 2,000 g centrifugation steps between washes discarding the supernatant). Thereafter, the pellet was re-suspended in buffer B (150 mM NaCl, HEPES pH7.5, 1 % (v/v) NP-40, 1 M Hexylene glycol, 1X Halt™ Protease Inhibitor Cocktail) and incubated for 30 min on ice. Samples were centrifuged for 10 min at 7,000 g at 4 °C and the supernatant (mitochondrial fraction) was transferred to a new 2 ml Eppendorf tube. To concentrate the samples, proteins were precipitated with 4x volumes of acetone for 1 h at – 20 °C and centrifugation for 1 h at 16,000 g at 4 °C. Protein pellets were re-suspended in 50 µl 1X LDS-loading buffer and boiled for 5 min at 95 °C.

### Cytoplasmic/nucleoplasmic vs chromatin fractionation

40-60 % confluent HEK293T cells on 10 cm plates were transfected with 3 µg expression construct (FLAG-FUS, FLAG-FUS PLD27YS or FLAG-FUS PLD27YS SV40NLS respectively) using Dogtor (OZ Biosciences) according to the manufacturer’s instructions. The medium was changed 24 h after transfection and cells were harvested 48 h after transfection using a cell scraper. Cells were transferred to a 15 ml Falcon tube and spun for 5 min at 200 g at 4 °C. After removal of the supernatant, cells were washed with 1 ml ice-cold PBS and transferred to a 1.5 ml Eppendorf tubed and pelleted for 3 min at 300 g at 4 °C. Cells were re-suspended in 1 ml buffer A (10 mM HEPES pH7.9, 10 mM KCl, 1.5 mM MgCl_2_, 0.34 M sucrose, 10 % glycerol, 0.1 % Triton-X-100, 1 mM DTT and 1X Halt™ Protease Inhibitor Cocktail) and incubated for 5 min on ice. The suspension was centrifuged for 4 min at 1’300 g at 4 °C and the supernatant (cytoplasmic fraction) was transferred to a new Eppendorf tube. The pellet was washed once with buffer A. Thereafter, the pellet was re-suspended in 1 ml buffer B (3 mM EDTA, 0.2 mM EGTA, 1 mM DTT and 1X Halt™ Protease Inhibitor Cocktail) and incubated for 5 min on ice. The suspension was centrifuged for 4 min at 1’700 g at 4 °C and the supernatant (nucleoplasmic fraction) was combined with the cytoplasmic fraction (forming the cytoplasmic/nucleoplasmic fraction). The pellet was washed once with buffer B. Then, the pellet was re-suspended in 1 ml buffer C (50 mM Tris pH7.5, 200 mM NaCl, 2 mM KCl, 1 mM EDTA, 0.5 % glycerol, 100 mM Urea and 1X Halt™ Protease Inhibitor Cocktail) and incubated in a heat block for 1 h at 37 °C at 1’400 rpm. After centrifugation for 5 min at 16’000 g, the supernatant (chromatin associated proteins) was transferred to a new Eppendorf tube. The pellet was subsequently washed with ultrapure water and buffer D (1 mM MgCl_2_, 1 mM CaCl_2_, 1X Halt™ Protease Inhibitor Cocktail). The pellet was then re-suspended in 884 µl buffer E (50 mM TRIS pH8.0, 10 mM NaCl, 1 mM MnSO_4_, 0.25 U/ml Cyanase (18542, Serva) and 1X Halt™ Protease Inhibitor Cocktail) and incubated for 15 min at 30 °C at 600 rpm. Thereafter, NaCl was increased to 600 mM through addition of 116 µl of 5 M NaCl. The samples was incubated for another 15 min at 37 °C at 1’400 rpm. After centrifugation for 5 min at 16’000 g, the supernatant (chromatin) was combined with the chromatin associated fraction (forming chromatin fraction). Fractions were supplemented with equal amounts of 2X LDS-loading buffer and boiled for 5 min at 95 °C.

### Immunofluorescence

HEK293T cells were grown on poly-D-lysine (Sigma Aldrich, A-003-E) coated 8-well slides (PEZGS0816, Milipore) for immunostaining experiments. HeLa cells were grown in uncoated 8-well slides. Cells were fixed for 20 minutes in 4 % PFA in PBS and subsequently washed 3 x with PBS. For permeabilization and blocking, cells were incubated for 45 minutes in 0.5 % Triton, 6 % BSA in TBS at room temperature. Primary antibodies were diluted in 0.1 % Triton, 6 % BSA in TBS (TBS +/+) and added to the cells overnight at 4 °C. Thereafter, cells were washed 3 x with TBS +/+ and incubated for 1 h at room temperature with secondary antibodies diluted in TBS +/+. Cells were counterstained with DAPI (100 ng/ml in PBS) for 10 min at room temperature. After two additional wash steps in PBS, cells were mounted with Vecashield Hardset mounting medium (H-1400, Vector Laboratories). If indicated, cells were stressed prior to fixation. Arsenic stress: 0.5 mM Sodium(meta)arsenite (NaAsO_2_) (S7400, Sigma Aldrich) for 1 h. Sorbitol stress: 0.4 M D-Sorbitol (S1876, Sigma Aldrich). Slides were analyzed using a Leica DMI6000 B microscope or a non-confocal Ti-E epifluorescence microscope, Nikon.

### Immunoblotting

Proteins from spinal cord of 9 month old wild type and heterozygous Δ14 mice were generated as previously described (Devoy et al., 2017). Protein lysates in LDS-loading buffer were separated on a NuPAGE 4-12% Bis Tris Midi Gel (WG1403A or WG1402BOX, Thermo Fisher) and transferred on a nitrocellulose membrane using the iBlot Gel Transfer System (Thermo Fisher) according to the manufacturer’s instructions. For analysis of SMRACA4 and SMARCA5, proteins were separated on a NuPAGE™ 3-8% Tris-Acetate Protein Gel (EA0375BOX, Thermo Fisher) and transferred on a nitrocellulose membrane using the iBlot™ 2 Gel Transfer Device (Thermo Fisher). Membranes were blocked in 2 % BSA in in 0.1% Tween in Tris-buffered saline (TBST). Membranes were incubated with primary antibodies diluted in TBST for 2 h at room temperature. After 5 x 5 min wash steps in TBST, membranes were incubated with secondary antibodies in TBST for 1 h at room temperature. For immunodetection of SMARCA4 and SMARCA5, the SuperSignal™ Western Blot Enhancer kit (46640, Thermo Fisher) was used according to the manufacturer’s instructions. To measure total protein levels from spinal cord lysates of mice, the REVERT™ Total Protein Stain Kit (926-11015, LI-COR) was used according to the manufacturer’s instructions. The washed membranes were analyzed and signal intensity was determined (if required for the respective experiment) using the Odyssey Infrared Imaging System (LI-COR). Statistical significance of immunoblotting results was determined by paired t-test.

### FUS autoregulation and SCN4A minigene reporter assay

To assess FUS autoregulation, 80 % confluent HeLa cells in 6wells were transfected with either 500 ng mock plasmid (control condition), pcDNA6F-FUS, pcDNA6F-FUS PLD27YS or pcDNA6F-FUS PLD27YS SV40NLS respectively. 24 h after transfection, cells were split into two 6wells. 72 h after transfection, cells were harvested (1 well into TRIZOL for RNA isolation, 1 well into 100 µl RIPA buffer containing 1X Halt™ Protease Inhibitor Cocktail. Before subsequent western blotting, the lysate incubated for 20 min on ice, spun for 15 min at 4 °C at 16’000 g and the supernatant was subsequently transferred to a new Eppendorf tube containing equal amount of 2X LDS-loading buffer and boiled for 5 min at 95 °C.

For the SCN4A minigene reporter assay, 80 % confluent HeLa cells in 6wells were transfected with 500 ng pSUPuro-scr or pSUPuro FUS for the CTR knockdown or the FUS knockdown, respectively. For the rescue condition, each 500 ng additional pcDNA6F-FUS, pcDNA6F-FUS PLD27YS or pcDNA6F-FUS PLD27YS SV40NLS were transfected. 24 h after transfection, the cells were split into two 6 wells and selection for the pSUPuro plasmids was started using 2 µg/ml Puromycin (CAS 58-58-2, Santa Cruz). Selection was maintained for 36 h. Cells were harvested 72 h after transfection as above (one well for RNA isolation, one well for western blotting).

### RT-qPCR

RNA was isolated from cells using TRIZOL (TRI Reagent™ Solution, AM9738, Thermo Fisher) supplemented with 1:100 β-Mercaptoethanol (A1108, PanReac Applichem) according to the manufacturer’s instructions. RNA for the SCN4A reporter assay was DNase treated using the TURBO DNA-free™ Kit (AM1907, Thermo Fisher) prior to cDNA synthesis according to the manufacturer’s instructions. RNA quantity was determined using the NanoDrop™ One/OneC Microvolume UV-Vis Spectrophotometer (Thermo Fisher). cDNA was generated from 1 µg RNA using the AffinityScript Multiple Temperature cDNA Synthesis Kit (00436, Agilent Technologies) according to the manufacturer’s instructions using random hexamer primers (150 ng/µl) (Sigma Aldrich). To confirm successful DNase treatment, a control reaction omitting the reverse transcription enzyme was prepared. The cDNA was diluted to a RNA concentration of 8 ng/µl. qPCR was performed using the Takyon No ROX SYBR 2X MasterMix blue dTTP (UF-NSMT-B0701, Eurogentec) with a final MgCl_2_ concentration of 4 mM. Samples were measured in duplicates: 3 µl of cDNA were amplified in a total volume of 15 µl containing each 600 nM forward and reverse primer using the Rotor-Gene Q 2plex Platform (Quiagen) with the following cycling parameters: 5 min 95 °C (initial denaturation), 20 secs 60 °C, 5 secs 95 °C (40 cycles). After cycling, a melting curve was recorded from 65 °C to 95 °C rising by 1 °C each step. Analysis was performed as described previously (Metze et al., 2013). Statistical significance of qPCR results was determined by unequal variances t-test.

### RNA FISH combined with immunofluorescence

HeLa cells were grown to 80% confluency in 6-well plates and transfected with 1 μg of the FUS expression constructs. The next day, 40,000 transfected cells were re-seeded into 8-well slides (Merck, PEZGS0816) and incubated overnight. FISH / IF was essentially performed as described in (Reber et al., 2016). In brief, the cells were fixed with 4% PFA for 15 minutes, permeabilized in 70% Ethanol at 4°C for 48 hours and blocked with blocking buffer (1% BSA (A7030, Sigma Aldrich) in PBS, supplemented with 2 mM Ribonucleoside Vanadyl Complexes (R3380, Sigma Aldrich). Antibodies were diluted in blocking buffer and incubated at room temperature for 1 hour (primary) and 2 hours (secondary), respectively. Subsequently, antibody complexes were cross-linked to their targets using 4 % PFA for 5 minutes. Following equilibration in 2x SSC (300 mM NaCl, 30 mM sodium citrate pH 7.0), and incubation in pre-hybridization buffer (15% Formamide (17899, Thermo Scientific), 10 mM sodium phosphate, 2 mM RVC in 2x SSC, pH 7.0) at 42°C for 10 minutes, 6-FAM azide labelled antisense probes were diluted to 0.5 ng/μl in hybridization buffer (15% Formamide, 10 mM sodium phosphate, 10% dextran sulfate (S4030, Merck), 0.2% BSA, 0.5 μg/μl E.coli tRNA, 0.5 μg/μl salmon sperm DNA (15632011, Invitrogen), 2 mM RVC in 2x SSC, pH 7.0) and hybridized to their targets over night at 42°C. The next day, unbound probes were removed by washing two times 30 minutes with pre-hybridization buffer and three times 10 minutes in high stringency wash solution (20 % Formamide, 2 mM RVC in 0.05 x SSC, pH 7.0). Then, the cells were washed three times with 2x SSC before mounting with aqueous Vectashield mounting medium containing DAPI (H-1200, Vectorlabs).

### Silver staining

Proteins in LDS-loading buffer were separated on a NuPAGE 4-12% Bis Tris Midi Gel (WG1403A, Thermo Fisher) or on a Bolt™ 4-12% Bis-Tris Plus Gel (NW04120BOX, Thermo Fisher). The gel was incubated for 2 h at room temperature in fixing solution (50 % MeOH, 12 % HAc, 0.05 % formalin). The gel was subsequently washed 3 x 20 min at room temperature in 35 % EtOH. Thereafter, the gel was sensitized in 0.02 % Na_2_S_2_O_3_ for 2 min, washed 3 x 5 min in ultrapure water and incubated for 20 min at room temperature in silver staining solution (0.2 % AgNO_3_, 0.076 % formalin). Then, the gel was washed 2 x 1 min in ultrapure water and developed in developing solution (6 % Na_2_CO_3_, 0.05 % formalin, 0.0004 % Na_2_S_2_O_3_). Upon desired band intensities, the reaction was stopped by replacing developing solution with stop solution (50 % MeOH, 12 % HAc) and incubating for 5 min.

## References

Alberti, S., and Carra, S. (2018). Quality Control of Membraneless Organelles. Journal of molecular biology 430, 4711–4729.

Alberti, S., and Dormann, D. (2019). Liquid-Liquid Phase Separation in Disease. Annual review of genetics.

Alberti, S., and Hyman Anthony, A. (2016). Are aberrant phase transitions a driver of cellular aging? BioEssays 38, 959–968.

Aulas, A., Stabile, S., and Vande Velde, C. (2012). Endogenous TDP-43, but not FUS, contributes to stress granule assembly via G3BP. Molecular Neurodegeneration 7, 1.

Baghirova, S., Hughes, B.G., Hendzel, M.J., and Schulz, R. (2015). Sequential fractionation and isolation of subcellular proteins from tissue or cultured cells. MethodsX 2, 440–445.

Bentmann, E., Neumann, M., Tahirovic, S., Rodde, R., Dormann, D., and Haass, C. (2012). Requirements for Stress Granule Recruitment of Fused in Sarcoma (FUS) and TAR DNA-binding Protein of 43 kDa (TDP-43). Journal of Biological Chemistry 287, 23079–23094.

Blokhuis, A.M., Koppers, M., Groen, E.J., van den Heuvel, D.M., Dini Modigliani, S., Anink, J.J., Fumoto, K., van Diggelen, F., Snelting, A., Sodaar, P., et al. (2016). Comparative interactomics analysis of different ALS-associated proteins identifies converging molecular pathways. Acta neuropathologica 132, 175–196.

Boehning, M., Dugast-Darzacq, C., Rankovic, M., Hansen, A.S., Yu, T., Marie-Nelly, H., McSwiggen, D.T., Kokic, G., Dailey, G.M., Cramer, P., et al. (2018). RNA polymerase II clustering through carboxy-terminal domain phase separation. Nat Struct Mol Biol 25, 833–840.

Boeynaems, S., Alberti, S., Fawzi, N.L., Mittag, T., Polymenidou, M., Rousseau, F., Schymkowitz, J., Shorter, J., Wolozin, B., Van Den Bosch, L., et al. (2018). Protein Phase Separation: A New Phase in Cell Biology. Trends Cell Biol 28, 420–435.

Boija, A., Klein, I.A., Sabari, B.R., Dall’Agnese, A., Coffey, E.L., Zamudio, A.V., Li, C.H., Shrinivas, K., Manteiga, J.C., Hannett, N.M., et al. (2018). Transcription Factors Activate Genes through the Phase-Separation Capacity of Their Activation Domains. Cell 175, 1842–1855 e1816.

Bosco, D.A., Lemay, N., Ko, H.K., Zhou, H., Burke, C., Kwiatkowski, J.T.J., Sapp, P., McKenna-Yasek, D., Brown, J.R.H., and Hayward, L.J. (2010). Mutant FUS proteins that cause amyotrophic lateral sclerosis incorporate into stress granules. Human molecular genetics 19, 4160–4175.

Brody, D.L., Jiang, H., Wildburger, N., and Esparza, T.J. (2017). Non-canonical soluble amyloid-beta aggregates and plaque buffering: controversies and future directions for target discovery in Alzheimer’s disease. Alzheimer’s research & therapy 9, 62.

Burke, Kathleen A., Janke, Abigail M., Rhine, Christy L., and Fawzi, Nicolas L. (2015). Residue-by-residue view of in vitro FUS granules that bind the C-terminal domain of RNA polymerase II. Molecular Cell 60, 231–241.

Cha, S.J., Choi, H.J., Kim, H.J., Choi, E.J., Song, K.H., Im, D.S., and Kim, K. (2019). Parkin expression reverses mitochondrial dysfunction in fused in sarcoma-induced amyotrophic lateral sclerosis. Insect molecular biology.

Chi, B., O’Connell, J.D., Yamazaki, T., Gangopadhyay, J., Gygi, S.P., and Reed, R. (2018). Interactome analyses revealed that the U1 snRNP machinery overlaps extensively with the RNAP II machinery and contains multiple ALS/SMA-causative proteins. Scientific reports 8, 8755.

Cho, W.K., Spille, J.H., Hecht, M., Lee, C., Li, C., Grube, V., and Cisse, II (2018). Mediator and RNA polymerase II clusters associate in transcription-dependent condensates. Science 361, 412–415.

Chong, S., Dugast-Darzacq, C., Liu, Z., Dong, P., Dailey, G.M., Cattoglio, C., Heckert, A., Banala, S., Lavis, L., Darzacq, X., et al. (2018). Imaging dynamic and selective low-complexity domain interactions that control gene transcription. Science 361.

Cleveland, D.W., and Rothstein, J.D. (2001). From Charcot to Lou Gehrig: deciphering selective motor neuron death in ALS. Nature reviews. Neuroscience 2, 806–819.

Coady, T.H., and Manley, J.L. (2015). ALS mutations in TLS/FUS disrupt target gene expression. Genes & development 29, 1696–1706.

Deng, J., Yang, M., Chen, Y., Chen, X., Liu, J., Sun, S., Cheng, H., Li, Y., Bigio, E.H., Mesulam, M., et al. (2015). FUS Interacts with HSP60 to Promote Mitochondrial Damage. PLoS genetics 11, e1005357.

Deshaies, J.-E., Shkreta, L., Moszczynski, A.J., Sidibé, H., Semmler, S., Fouillen, A., Bennett, E.R., Bekenstein, U., Destroismaisons, L., Toutant, J., et al. (2018). TDP-43 regulates the alternative splicing of hnRNP A1 to yield an aggregation-prone variant in amyotrophic lateral sclerosis. Brain : a journal of neurology 141, 1320–1333.

Devoy, A., Kalmar, B., Stewart, M., Park, H., Burke, B., Noy, S.J., Redhead, Y., Humphrey, J., Lo, K., Jaeger, J., et al. (2017). Humanized mutant FUS drives progressive motor neuron degeneration without aggregation in ‘FUSDelta14’ knockin mice. Brain : a journal of neurology 140, 2797–2805.

Dormann, D., Rodde, R., Edbauer, D., Bentmann, E., Fischer, I., Hruscha, A., Than, M.E., Mackenzie, I.R.A., Capell, A., Schmid, B., et al. (2010). ALS-associated fused in sarcoma (FUS) mutations disrupt Transportin-mediated nuclear import. The EMBO journal 29, 2841–2857.

Ederle, H., Funk, C., Abou-Ajram, C., Hutten, S., Funk, E.B.E., Kehlenbach, R.H., Bailer, S.M., and Dormann, D. (2018). Nuclear egress of TDP-43 and FUS occurs independently of Exportin-1/CRM1. Scientific reports 8, 7084.

Fridy, P.C., Li, Y., Keegan, S., Thompson, M.K., Nudelman, I., Scheid, J.F., Oeffinger, M., Nussenzweig, M.C., Fenyo, D., Chait, B.T., et al. (2014). A robust pipeline for rapid production of versatile nanobody repertoires. Nature methods 11, 1253–1260.

Gerbino, V., Carrì, M.T., Cozzolino, M., and Achsel, T. (2013). Mislocalised FUS mutants stall spliceosomal snRNPs in the cytoplasm. Neurobiology of Disease 55, 120–128.

Groen, E.J.N., Fumoto, K., Blokhuis, A.M., Engelen-Lee, J., Zhou, Y., van den Heuvel, D.M.A., Koppers, M., van Diggelen, F., van Heest, J., Demmers, J.A.A., et al. (2013). ALS-associated mutations in FUS disrupt the axonal distribution and function of SMN. Human molecular genetics 22, 3690–3704.

Gunasekera, K., Wuthrich, D., Braga-Lagache, S., Heller, M., and Ochsenreiter, T. (2012). Proteome remodelling during development from blood to insect-form Trypanosoma brucei quantified by SILAC and mass spectrometry. BMC Genomics 13, 556.

Hein, Marco Y., Hubner, Nina C., Poser, I., Cox, J., Nagaraj, N., Toyoda, Y., Gak, Igor A., Weisswange, I., Mansfeld, J., Buchholz, F., et al. (2015). A Human Interactome in Three Quantitative Dimensions Organized by Stoichiometries and Abundances. Cell 163, 712–723.

Hock, E.M., Maniecka, Z., Hruska-Plochan, M., Reber, S., Laferriere, F., Sahadevan, M.K.S., Ederle, H., Gittings, L., Pelkmans, L., Dupuis, L., et al. (2018). Hypertonic Stress Causes Cytoplasmic Translocation of Neuronal, but Not Astrocytic, FUS due to Impaired Transportin Function. Cell reports 24, 987–1000 e1007.

Hofweber, M., Hutten, S., Bourgeois, B., Spreitzer, E., Niedner-Boblenz, A., Schifferer, M., Ruepp, M.-D., Simons, M., Niessing, D., Madl, T., et al. (2018). Phase Separation of FUS Is Suppressed by Its Nuclear Import Receptor and Arginine Methylation. Cell 173, 706–719.e713.

Hyman, A.A., Weber, C.A., and Julicher, F. (2014). Liquid-liquid phase separation in biology. Annual review of cell and developmental biology 30, 39–58.

Jutzi, D., Akinyi, M.V., Mechtersheimer, J., Frilander, M.J., and Ruepp, M.-D. (2018). The emerging role of minor intron splicing in neurological disorders. Cell Stress 2, 40–54.

Kamelgarn, M., Chen, J., Kuang, L., Arenas, A., Zhai, J., Zhu, H., and Gal, J. (2016). Proteomic analysis of FUS interacting proteins provides insights into FUS function and its role in ALS. Biochimica et Biophysica Acta (BBA) - Molecular Basis of Disease 1862, 2004–2014.

Kang, J., Lim, L., Lu, Y., and Song, J. (2019). A unified mechanism for LLPS of ALS/FTLD-causing FUS as well as its modulation by ATP and oligonucleic acids. PLoS biology 17, e3000327.

Kato, M., Han, Tina W., Xie, S., Shi, K., Du, X., Wu, Leeju C., Mirzaei, H., Goldsmith, Elizabeth J., Longgood, J., Pei, J., et al. (2012). Cell-free Formation of RNA Granules: Low Complexity Sequence Domains Form Dynamic Fibers within Hydrogels. Cell 149, 753–767.

Krey, J.F., Wilmarth, P.A., Shin, J.B., Klimek, J., Sherman, N.E., Jeffery, E.D., Choi, D., David, L.L., and Barr-Gillespie, P.G. (2014). Accurate label-free protein quantitation with high- and low-resolution mass spectrometers. Journal of proteome research 13, 1034–1044.

Krueger, F. (2015). A wrapper tool around Cutadapt and FastQC to consistently apply quality and adapter trimming to FastQ files.

Kwiatkowski, T.J., Jr., Bosco, D.A., Leclerc, A.L., Tamrazian, E., Vanderburg, C.R., Russ, C., Davis, A., Gilchrist, J., Kasarskis, E.J., Munsat, T., et al. (2009). Mutations in the FUS/TLS gene on chromosome 16 cause familial amyotrophic lateral sclerosis. Science 323, 1205–1208.

Liao, Y., Wang, J., Jaehnig, E.J., Shi, Z., and Zhang, B. (2019). WebGestalt 2019: gene set analysis toolkit with revamped UIs and APIs. Nucleic acids research.

Linden, M., Thomsen, C., Grundevik, P., Jonasson, E., Andersson, D., Runnberg, R., Dolatabadi, S., Vannas, C., Luna Santamariotaa, M., Fagman, H., et al. (2019). FET family fusion oncoproteins target the SWI/SNF chromatin remodeling complex. EMBO reports 20.

Lindstrom, M., and Liu, B. (2018). Yeast as a Model to Unravel Mechanisms Behind FUS Toxicity in Amyotrophic Lateral Sclerosis. Front Mol Neurosci 11, 218.

Ling, S.C., Polymenidou, M., and Cleveland, D.W. (2013). Converging mechanisms in ALS and FTD: disrupted RNA and protein homeostasis. Neuron 79, 416–438.

Loughlin, F.E., Lukavsky, P.J., Kazeeva, T., Reber, S., Hock, E.M., Colombo, M., Von Schroetter, C., Pauli, P., Clery, A., Muhlemann, O., et al. (2019). The Solution Structure of FUS Bound to RNA Reveals a Bipartite Mode of RNA Recognition with Both Sequence and Shape Specificity. Mol Cell 73, 490–504 e496.

Martin, M. (2011). Cutadapt removes adapter sequences from high-throughput sequencing reads. EMBnet. journal 17.

Matus, S., Bosco, D.A., and Hetz, C. (2014). Autophagy meets fused in sarcoma-positive stress granules. Neurobiology of aging 35, 2832–2835.

Meissner, M., Lopato, S., Gotzmann, J., Sauermann, G., and Barta, A. (2003). Proto-oncoprotein tls/fus is associated to the nuclear matrix and complexed with splicing factors ptb, srm160, and sr proteins. Experimental Cell Research 283, 184–195.

Metze, S., Herzog, V.A., Ruepp, M.D., and Muhlemann, O. (2013). Comparison of EJC-enhanced and EJC-independent NMD in human cells reveals two partially redundant degradation pathways. RNA 19, 1432–1448.

Mitchell, J.C., McGoldrick, P., Vance, C., Hortobagyi, T., Sreedharan, J., Rogelj, B., Tudor, E.L., Smith, B.N., Klasen, C., Miller, C.C.J., et al. (2013). Overexpression of human wild-type FUS causes progressive motor neuron degeneration in an age- and dose-dependent fashion. Acta neuropathologica 125, 273–288.

Monahan, Z., Ryan, V.H., Janke, A.M., Burke, K.A., Rhoads, S.N., Zerze, G.H., Meally, R., Dignon, G.L., Conicella, A.E., Zheng, W., et al. (2017). Phosphorylation of the FUS low-complexity domain disrupts phase separation, aggregation, and toxicity. The EMBO journal 36, 2951–2967.

Murray, D.T., Kato, M., Lin, Y., Thurber, K.R., Hung, I., McKnight, S.L., and Tycko, R. (2017). Structure of FUS Protein Fibrils and Its Relevance to Self-Assembly and Phase Separation of Low-Complexity Domains. Cell 171, 615–627.e616.

Patel, A., Lee, Hyun O., Jawerth, L., Maharana, S., Jahnel, M., Hein, Marco Y., Stoynov, S., Mahamid, J., Saha, S., Franzmann, Titus M., et al. (2015). A Liquid-to-Solid Phase Transition of the ALS Protein FUS Accelerated by Disease Mutation. Cell 162, 1066–1077.

Patel, A.A., McCarthy, M., and Steitz, J.A. (2002). The splicing of U12-type introns can be a rate-limiting step in gene expression. The EMBO journal 21, 3804–3815.

Patro, R., Duggal, G., Love, M.I., Irizarry, R.A., and Kingsford, C. (2017). Salmon provides fast and bias-aware quantification of transcript expression. Nature methods 14, 417–419.

Prasad, D.D., Ouchida, M., Lee, L., Rao, V.N., and Reddy, E.S. (1994). TLS/FUS fusion domain of TLS/FUS-erg chimeric protein resulting from the t(16;21) chromosomal translocation in human myeloid leukemia functions as a transcriptional activation domain. Oncogene 9, 3717–3729.

Qamar, S., Wang, G., Randle, S.J., Ruggeri, F.S., Varela, J.A., Lin, J.Q., Phillips, E.C., Miyashita, A., Williams, D., Strohl, F., et al. (2018). FUS Phase Separation Is Modulated by a Molecular Chaperone and Methylation of Arginine Cation-pi Interactions. Cell 173, 720–734 e715.

Raczynska, K.D., Ruepp, M.D., Brzek, A., Reber, S., Romeo, V., Rindlisbacher, B., Heller, M., Szweykowska-Kulinska, Z., Jarmolowski, A., and Schumperli, D. (2015). FUS/TLS contributes to replication-dependent histone gene expression by interaction with U7 snRNPs and histone-specific transcription factors. Nucleic acids research 43, 9711–9728.

Reber, S., Stettler, J., Filosa, G., Colombo, M., Jutzi, D., Lenzken, S.C., Schweingruber, C., Bruggmann, R., Bachi, A., Barabino, S.M., et al. (2016). Minor intron splicing is regulated by FUS and affected by ALS-associated FUS mutants. The EMBO journal 35, 1504–1521.

Robinson, M.D., McCarthy, D.J., and Smyth, G.K. (2010). edgeR: a Bioconductor package for differential expression analysis of digital gene expression data. Bioinformatics 26, 139–140.

Sabari, B.R., Dall’Agnese, A., Boija, A., Klein, I.A., Coffey, E.L., Shrinivas, K., Abraham, B.J., Hannett, N.M., Zamudio, A.V., Manteiga, J.C., et al. (2018). Coactivator condensation at super-enhancers links phase separation and gene control. Science 361.

Sabatelli, M., Moncada, A., Conte, A., Lattante, S., Marangi, G., Luigetti, M., Lucchini, M., Mirabella, M., Romano, A., Del Grande, A., et al. (2013). Mutations in the 3ʹ untranslated region of FUS causing FUS overexpression are associated with amyotrophic lateral sclerosis. Human molecular genetics 22, 4748–4755.

Scekic-Zahirovic, J., Sendscheid, O., El Oussini, H., Jambeau, M., Sun, Y., Mersmann, S., Wagner, M., Dieterlé, S., Sinniger, J., Dirrig-Grosch, S., et al. (2016). Toxic gain of function from mutant FUS protein is crucial to trigger cell autonomous motor neuron loss. The EMBO journal 35, 1077–1097.

Schmidt, H.B., Barreau, A., and Rohatgi, R. (2019). Decoding and recoding phase behavior of TDP43 reveals that phase separation is not required for splicing function. bioRxiv, 548339.

Schwartz, J.C., Ebmeier, C.C., Podell, E.R., Heimiller, J., Taatjes, D.J., and Cech, T.R. (2012). FUS binds the CTD of RNA polymerase II and regulates its phosphorylation at Ser2. Genes & development 26, 2690–2695.

Schwartz, Jacob C., Wang, X., Podell, Elaine R., and Cech, Thomas R. (2013). RNA Seeds Higher-Order Assembly of FUS Protein. Cell reports 5, 918–925.

Sharma, A., Lyashchenko, A.K., Lu, L., Nasrabady, S.E., Elmaleh, M., Mendelsohn, M., Nemes, A., Tapia, J.C., Mentis, G.Z., and Shneider, N.A. (2016). ALS-associated mutant FUS induces selective motor neuron degeneration through toxic gain of function. Nature communications 7, 10465.

Shelkovnikova, T.A., Robinson, H.K., Troakes, C., Ninkina, N., and Buchman, V.L. (2014). Compromised paraspeckle formation as a pathogenic factor in FUSopathies. Human molecular genetics 23, 2298–2312.

Singh, J., and Padgett, R.A. (2009). Rates of in situ transcription and splicing in large human genes. Nat Struct Mol Biol 16, 1128–1133.

Soneson, C., Love, M.I., and Robinson, M.D. (2015). Differential analyses for RNA-seq: transcript-level estimates improve gene-level inferences. F1000Research 4, 1521.

Steyaert, J., Scheveneels, W., Vanneste, J., Van Damme, P., Robberecht, W., Callaerts, P., Bogaert, E., and Van Den Bosch, L. (2018). FUS-induced neurotoxicity in Drosophila is prevented by downregulating nucleocytoplasmic transport proteins. Human molecular genetics 27, 4103–4116.

Stoica, R., Paillusson, S., Gomez-Suaga, P., Mitchell, J.C., Lau, D.H.W., Gray, E.H., Sancho, R.M., Vizcay-Barrena, G., De Vos, K.J., Shaw, C.E., et al. (2016). ALS/FTD-associated FUS activates GSK-3β to disrupt the VAPB–PTPIP51 interaction and ER–mitochondria associations. EMBO reports 17, 1326–1342.

Sun, S., Ling, S.-C., Qiu, J., Albuquerque, C.P., Zhou, Y., Tokunaga, S., Li, H., Qiu, H., Bui, A., Yeo, G.W., et al. (2015). ALS-causative mutations in FUS/TLS confer gain and loss of function by altered association with SMN and U1-snRNP. Nature communications 6, 6171.

Szklarczyk, D., Franceschini, A., Wyder, S., Forslund, K., Heller, D., Huerta-Cepas, J., Simonovic, M., Roth, A., Santos, A., Tsafou, K.P., et al. (2015). STRING v10: protein-protein interaction networks, integrated over the tree of life. Nucleic acids research 43, D447–452.

Tang, L. (2019). Liquid phase separation. Nature methods 16, 18.

Thompson, V.F., Victor, R.A., Morera, A.A., Moinpour, M., Liu, M.N., Kisiel, C.C., Pickrel, K., Springhower, C.E., and Schwartz, J.C. (2018). Transcription-Dependent Formation of Nuclear Granules Containing FUS and RNA Pol II. Biochemistry 57, 7021–7032.

Vance, C., Rogelj, B., Hortobagyi, T., De Vos, K.J., Nishimura, A.L., Sreedharan, J., Hu, X., Smith, B., Ruddy, D., Wright, P., et al. (2009). Mutations in FUS, an RNA processing protein, cause familial amyotrophic lateral sclerosis type 6. Science 323, 1208–1211.

Wang, C., and Youle, R.J. (2009). The role of mitochondria in apoptosis. Annual review of genetics 43, 95–118.

Wang, J., Choi, J.M., Holehouse, A.S., Lee, H.O., Zhang, X., Jahnel, M., Maharana, S., Lemaitre, R., Pozniakovsky, A., Drechsel, D., et al. (2018). A Molecular Grammar Governing the Driving Forces for Phase Separation of Prion-like RNA Binding Proteins. Cell 174, 688–699 e616.

Wang, T., Jiang, X., Chen, G., and Xu, J. (2015). Interaction of amyotrophic lateral sclerosis/frontotemporal lobar degeneration-associated fused-in-sarcoma with proteins involved in metabolic and protein degradation pathways. Neurobiology of aging 36, 527–535.

Wolozin, B., and Ivanov, P. (2019). Stress granules and neurodegeneration. Nature reviews. Neuroscience.

Yang, L., Gal, J., Chen, J., and Zhu, H. (2014). Self-assembled FUS binds active chromatin and regulates gene transcription. Proceedings of the National Academy of Sciences 111, 17809–17814.

Younis, I., Dittmar, K., Wang, W., Foley, S.W., Berg, M.G., Hu, K.Y., Wei, Z., Wan, L., and Dreyfuss, G. (2013). Minor introns are embedded molecular switches regulated by highly unstable U6atac snRNA. eLife 2, e00780.

Yu, S., Jordan-Pla, A., Ganez-Zapater, A., Jain, S., Rolicka, A., Ostlund Farrants, A.K., and Visa, N. (2018). SWI/SNF interacts with cleavage and polyadenylation factors and facilitates pre-mRNA 3’ end processing. Nucleic acids research 46, 8557–8573.

Yu, Y., and Reed, R. (2015). FUS functions in coupling transcription to splicing by mediating an interaction between RNAP II and U1 snRNP. Proceedings of the National Academy of Sciences 112, 8608–8613.

Zhang, T., Wu, Y.-C., Mullane, P., Ji, Y.J., Liu, H., He, L., Arora, A., Hwang, H.-Y., Alessi, A.F., Niaki, A.G., et al. (2018). FUS Regulates Activity of MicroRNA-Mediated Gene Silencing. Molecular Cell 69, 787–801.e788.

Zinszner, H., Albalat, R., and Ron, D. (1994). A novel effector domain from the RNA-binding protein TLS or EWS is required for oncogenic transformation by CHOP. Genes & development 8, 2513–2526.

