## Supplemental Material for "The phase separation-dependent FUS interactome reveals nuclear and cytoplasmic function of liquid-liquid phase separation"

### Supplemental Material and methods

#### *Plasmids*

pcDNA6F-FUS, pcDNA6F-FUS P525L, pcDNA3-FLAG-eGFP, pSUPuro-scr, pSUPuro-FUS and the SCN4A reporter minigene are described elsewhere(Reber et al., 2016; Rufener and Muhlemann, 2013). To generate pcDNA6F-FUS PLD27YS, a gene synthesis (GeneArt, Life Technologies) coding for the N-terminal domain of FUS with 27 tyrosines substituted with serines according to(Kato et al., 2012) flanked by XhoI and EcoRI restriction sites was used to clone the FUS N-term (PLD27YS) into the XhoI, EcoRI restriction sites of pcDNA6F-FUS (thereby replacing the wild type FUS N-term). To clone pcDNA6F-FUS PLD27YS P525L the aforementioned strategy was used and the FUS N-term (PLD27YS) was cloned into the XhoI, EcoRI restriction sites of pcDNA6F-FUS P525L. To generate pcDNA6F-FUS PLD27YS SV40NLS, the sequence coding for FUS PLD27YS was PCR amplified from pcDNA6F-FUS PLD27YS using the CloneAmp HiFi PCR Premix (639298, Clontech) with the forward primer 5’-ATAGGGAGACCCAAGCTGGCTAG-3’ and the reverse primer introducing the SV40NLS (SV40 nuclear localization signal) 5’-GATTGGGCCCTTCACTTGTCCTCCACTTTGCGTTTCTTTTTGGGATACGGCCTCTCCCTGCGATCC-3’. The PCR product was digested with XhoI and ApaI and cloned into the XhoI, ApaI site of pcDNA6f-FUS (replacing the coding sequence for FUS). To generate pcDNA6F-GFP-GSG15-FUS and pcDNA6F-eGFP-GSG15-FUS P525L, the sequence coding for the N-terminal FLAG tag was removed from pcDNA6F-FUS and pcDNA6F-FUS P525L, respectively and was replaced with the coding sequence for eGFP followed by a GSG15 linker which was ordered as a gene synthesis (General Biosystems) and cloned into the XbaI, XhoI sites.

#### *qPCR Primers*

| Primer | 5’-sequence-3’ |
|---|---|
| qPCR endogenous FUS fwd | AGCGGTGTTGGAACTTCG |
| qPCR endogenous FUS rev | GACTGCTCTGCTGGGAATAG |
| qPCR β-actin fwd | TCCATCATGAAGTGTGACGT |
| qPCR β-actin rev | TACTCCTGCTTGCTGATCCAC |
| qPCR SCN4A total fwd | CAAGGGCAAGGCCATCTTC |
| qPCR SCN4A total rev | GCATGGATGAGCACCTTGATG |
| qPCR SCN4A spliced fwd | ACAAGGGCAAGGCCATCTTC |
| qPCR SCN4A unspliced rev | CATGCTGAACAGCGCATGG |

#### *Antibodies*

The polyclonal rabbit anti-FUS antibody, the Y12 monoclonal antibody, the rabbit anti-CPSF6 (also CFIm68) and rabbit anti-NUDT21 (also CFIm25) are described elsewhere(Lerner et al., 1981; Raczynska et al., 2015; Ruegsegger et al., 1998). Additional antibodies used for this study: mouse anti-FUS (4H11) (sc-47711, Santa Cruz), mouse anti-FLAG M2 antibody (F1804, Sigma-Aldrich), rabbit anti-FLAG

(14793S, Cell Signaling Technology), mouse anti-tyrosine tubulin (T9028, Sigma-Aldrich), mouse anti-VDAC1 (ab14734, abcam), mouse anti-TOM20 (sc-17764, Santa Cruz), rabbit anti-PARP1 (ab227244, abcam), rabbit anti-Lig3 (ab125434, abcam), rabbit anti-hnRNPA2B1 (ab31645, abcam), mouse anti-cytochrome c (sc-13156, Santa Cruz), mouse anti-GAPDH (sc-32233, Santa Cruz), mouse anti-hSNF2H (also SMARCA5) (sc-365727, Santa Cruz), mouse anti-Brg-1 (also SMARCA4) (sc-17796, Santa Cruz), rabbit anti-hnRNPH (A300-511A, Bethyl Laboratories), mouse anti-hnRNPA1 (sc-56700, Santa Cruz), mouse anti-SNRPC (5C9) (also U1C) (sc-101548, Santa Cruz), mouse anti-SNRPA SNRPA (BJ-7) (also U1A) (sc-101149, Santa Cruz), rabbit anti-GFP (ab6556, abcam), goat anti-GFP (AB0020-200, SICGEN), rabbit anti-histone H3 (4499, Cell Signaling Technology), mouse anti-RNAPII (CTD4H8) (05-623B, Milipore), mouse anti-G3BP (611126, BD Transduction Laboratories), goat anti-TIAR (sc-1749, Santa Cruz), donkey anti-goat IRDye800CW (926-32214, LI-COR Biosciences), goat anti-mouse IRDye800CW (925-32210, LI-COR), goat anti-rabbit IRDye800CW (926-32211, LI-COR), goat anti-mouse IRDye680LT (926-68020, LI-COR), goat anti-rabbit IRDye680LT (926-68021, LI-COR), donkey anti-mouse IRDye680LT (926-68022, LI-COR Biosciences), donkey anti-rabbit AF488 (R37118, Thermo Fisher), donkey anti-goat AF568 (A-11057, Thermo Fisher), goat anti-rabbit AF488 (A27034, Thermo Fisher), goat anti-mouse AF546 (A-11003, Thermo Fisher), donkey anti-rabbit AF-546 (A10040, Thermo Fisher), donkey anti-mouse AF-488 (A21202, Thermo Fisher).

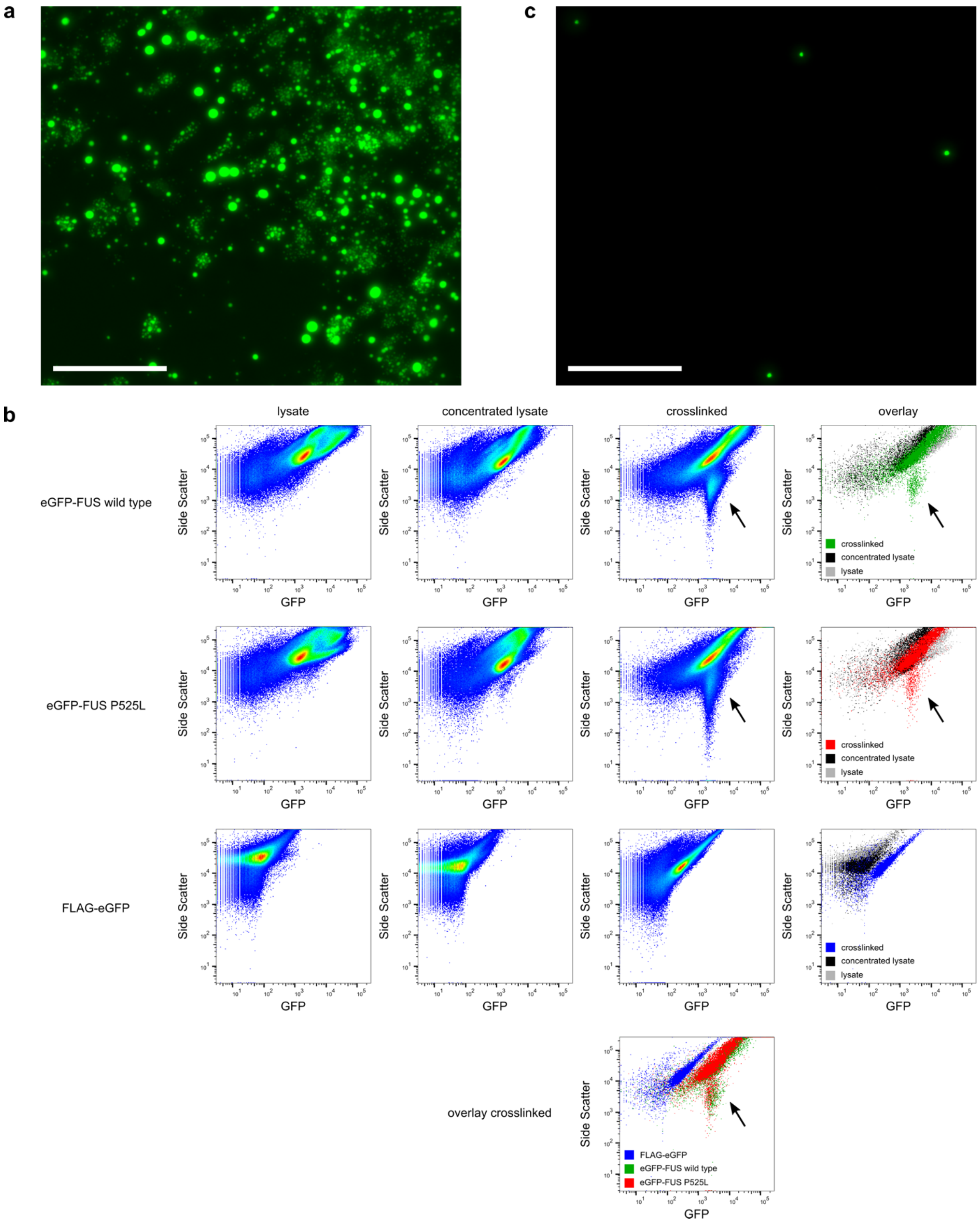


*Supplementary Figure 1 related to Figure 1*

Analysis of eGFP-FUS droplet formation by fluorescence microscopy and flow cytometry. **a** Picture of the concentrated cell lysate before crosslinking. A large number of eGFP-FUS droplets of varying size and cell debris are visible. Scale bar = 75 µm. **b** Flow cytometric analysis of lysates (first column), concentrated lysates (second column) and crosslinked lysates (third column) from cells which were expressing eGFP-FUS (top row), eGFP-FUS P525L (second row) or FLAG-eGFP (third row). The last column shows the overlay of each row. The fourth row shows the overlay of the third column. The population of droplets, which are only stable after crosslinking, are indicated with black arrows. **c** Picture of purified droplets after sorting. Scale bar = 75 µm.

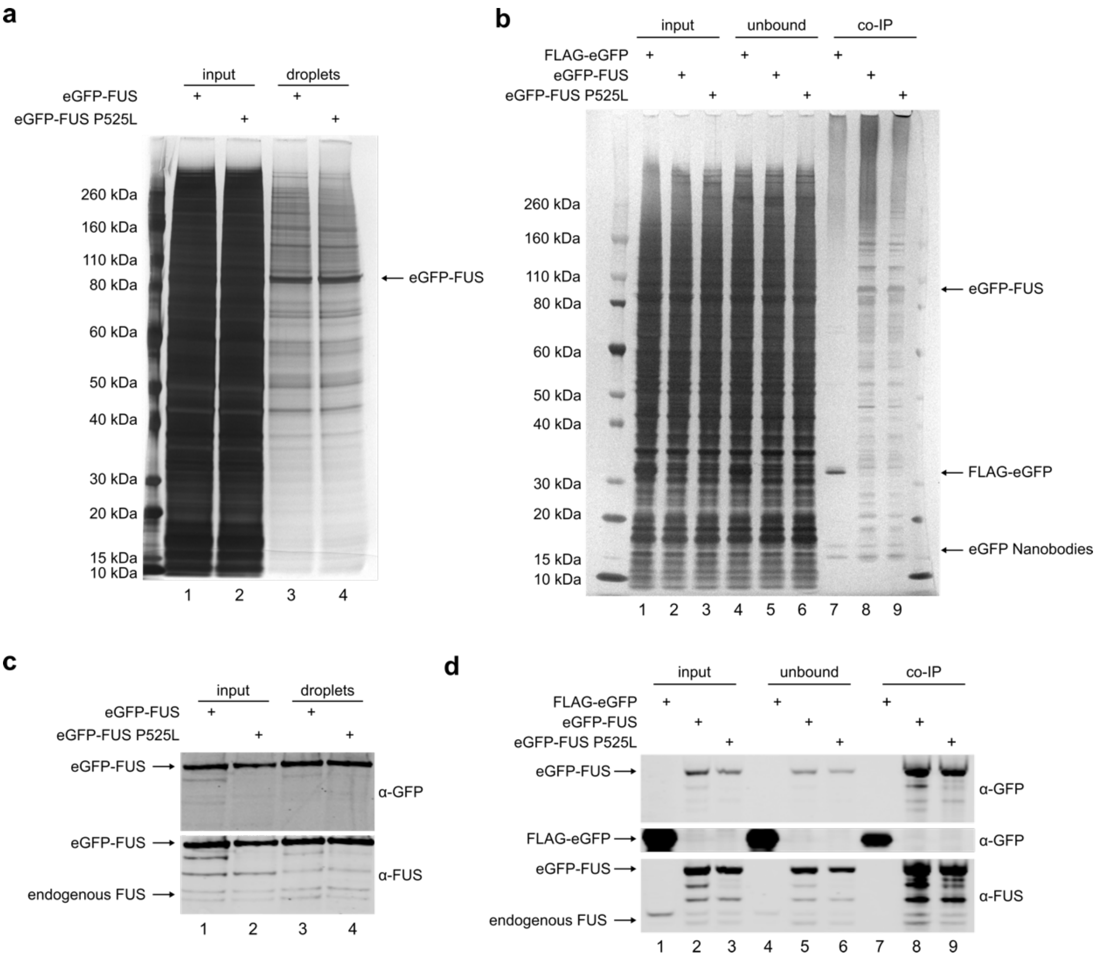


*Supplementary Figure 2 related to Figure 1*

Silver stainings and western blots of droplet purification and co-immunoprecipitation experiments separated on 4-12 % Bis-Tris gels. **a** Silver staining of input (lane 1-2) and purified droplets (lane 3-4). Arrow indicates eGFP-FUS and eGFP-FUS P525L, respectively. **b** Silver staining of input (lane 1-3), unbound fractions (4-6) and co-immunoprecipitated proteins together with FLAG-eGFP (lane 7), eGFP-FUS (lane 8) and eGFP-FUS P525L (lane 9). Arrows indicate baits and eGFP Nanobodies which were partially boiled off the beads (2 bands at > 15 kDa). **c** Western blot of samples shown in a. Proteins were transferred on a nitrocellulose membrane and the membrane was subsequently probed with anti-GFP (top panel) and anti-FUS (bottom panel) antibodies. **d** Western blot of samples shown in b. The nitrocellulose membrane was treated as in c.

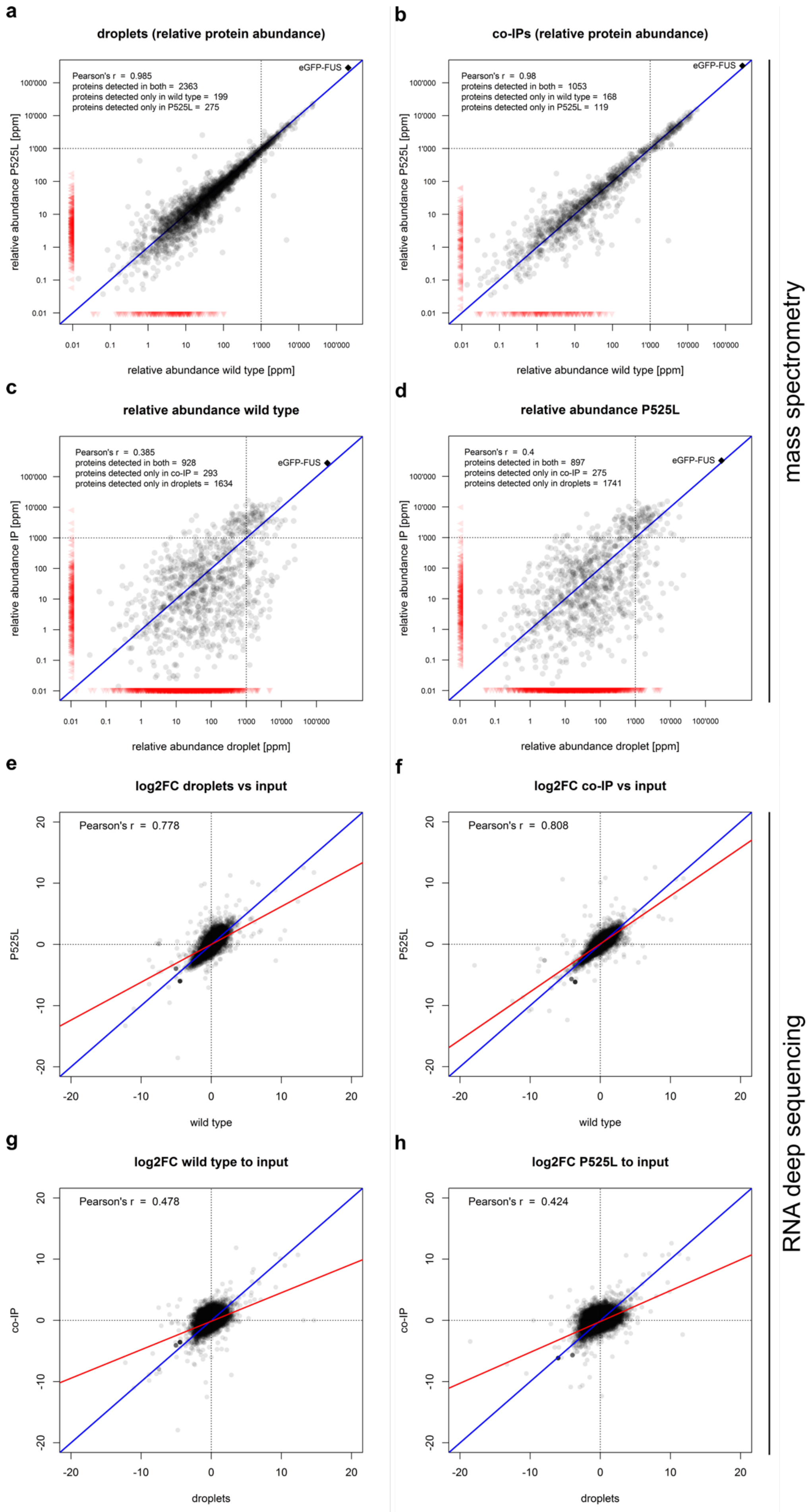
a
droplets (relative protein abundance)
Pearson's r = 0.985
proteins detected in both = 2363
proteins detected only in wild type = 199
proteins detected only in P525L = 275
eGFP-FUS
relative abundance P525L [ppm]
relative abundance wild type [ppm]
b
co-IPs (relative protein abundance)
Pearson's r = 0.98
proteins detected in both = 1053
proteins detected only in wild type = 168
proteins detected only in P525L = 119
eGFP-FUS
relative abundance P525L [ppm]
relative abundance wild type [ppm]
c
relative abundance wild type
Pearson's r = 0.385
proteins detected in both = 928
proteins detected only in co-IP = 293
proteins detected only in droplets = 1634
eGFP-FUS
relative abundance IP [ppm]
relative abundance droplet [ppm]
d
relative abundance P525L
Pearson's r = 0.4
proteins detected in both = 897
proteins detected only in co-IP = 275
proteins detected only in droplets = 1741
eGFP-FUS
relative abundance IP [ppm]
relative abundance droplet [ppm]
mass spectrometry
e
log2FC droplets vs input
Pearson's r = 0.778
P525L
wild type
f
log2FC co-IP vs input
Pearson's r = 0.808
P525L
wild type
g
log2FC wild type to input
Pearson's r = 0.478
co-IP
droplets
h
log2FC P525L to input
Pearson's r = 0.424
co-IP
droplets
RNA deep sequencing

*Supplementary Figure 3 related to Figure 1*

Plots showing correlations between proteins (a-d) and RNAs (e-h) detected in quantitative mass spectrometry and RNAdeep sequencing, respectively. Note that there is a high correlation between wild type FUS and P525L FUS samples within the same experiment (a-b and e-f) while a much lower or no correlation can be observed if droplet and co-IP samples are compared (c-d and g-h). **a** Plot showing relative abundance in parts per million (ppm) of all proteins which were detected in wild type (x-axis) and P525L FUS (y-axis) droplets. The dotted lines label 1,000 ppm (= 0.1 %). The blue line depicts perfect correlation. Black dots represent proteins that were detected in both samples. Red triangles represent proteins that were detected in one sample only. The bait (eGFP-FUS) is represented as a black square. Pearson's correlation coefficient was calculated on the non-logarithmic values and the bait (eGFP-FUS) was excluded from the calculation. **b** Same plot as in a, but showing relative abundances of proteins detected in the co-IP experiments. **c** Same plot as in a, but comparing droplet and co-IP experiment of wild type FUS samples. **d** Same as in a, but comparing droplet and co-IP experiment of P525L FUS samples. **e** Plot showing the log2 fold change of RNA isolated from droplets compared to the input together with wild type (x-axis) and P525L FUS (y-axis). The blue line depicts perfect correlation, the red line the actual correlation between the samples. Each black dot represents one gene (n = 13,096) **f** Same as in e, but showing log2 fold changes of RNAs isolated in the co-IP experiment. **g** Same plot as in e, but comparing droplet and co-IP experiments of wild type FUS samples. **h** Same as in a, but comparing droplet and co-IP experiment of P525L FUS samples.

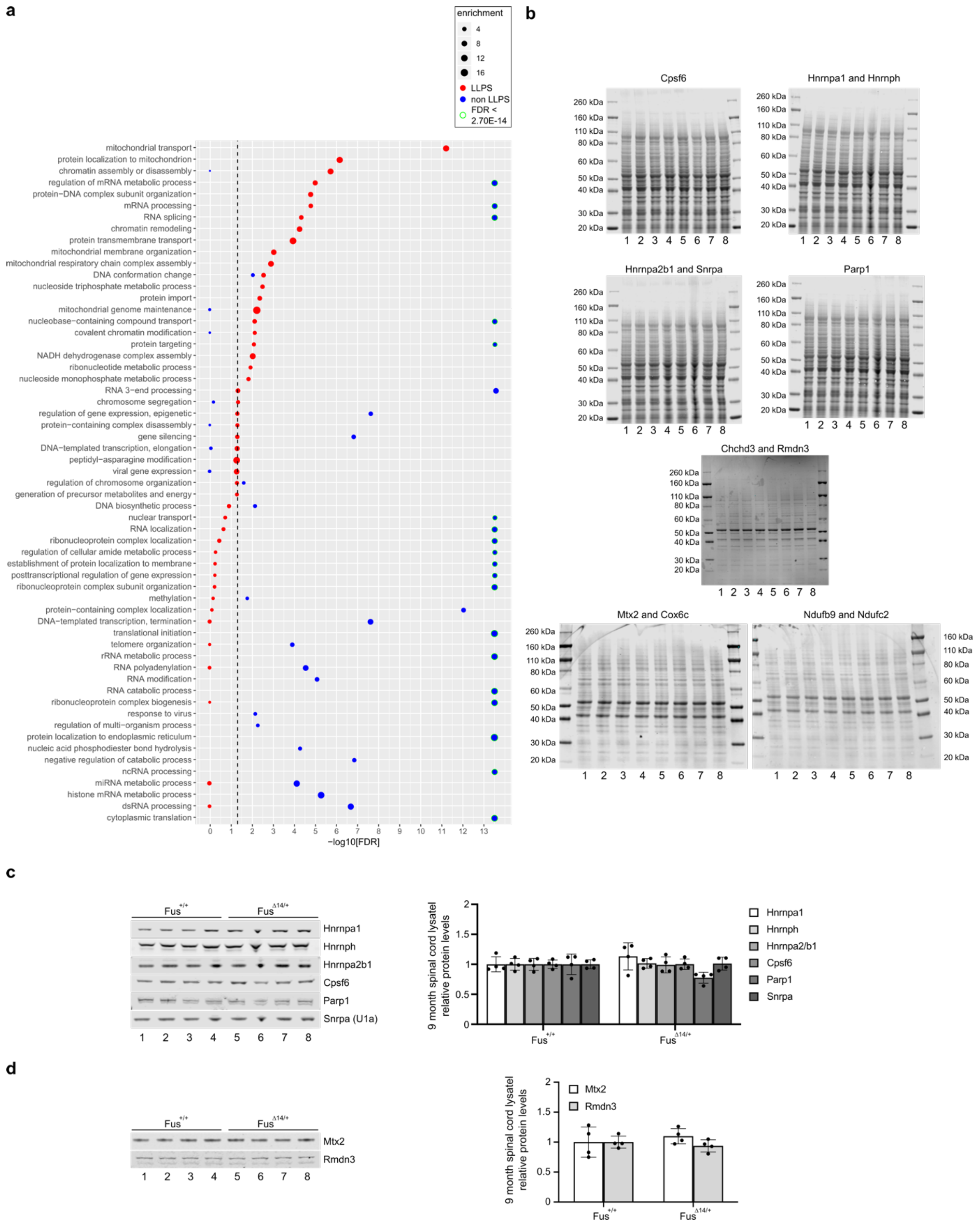

*Supplementary Figure 4 related to Figure 2*

A GO term analysis of LLPS-specific FUS interactors (red) and proteins interacting with FUS under non-LLPS conditions (blue). The dotted line indicates a FDR < 0.05. All significantly enriched GO terms for both FUS interactomes are shown. Missing dots indicate that the respective GO term was not detected in the respective FUS interactome. **b** Total protein stainings used for quantification of protein levels and normalization in Figure 2g and Supplementary Figure 4c and d. Note that while membranes were routinely blocked after total protein staining (and scan) were performed, the total protein staining for Chchd3 and Rmdn3 was performed after the membrane was blocked. **c** Western blot (left) and quantification (right) of proteins involved in RNA splicing and chromatin remodelling. **d** Western blot (left) and quantification (right) of proteins with mitochondrial function.

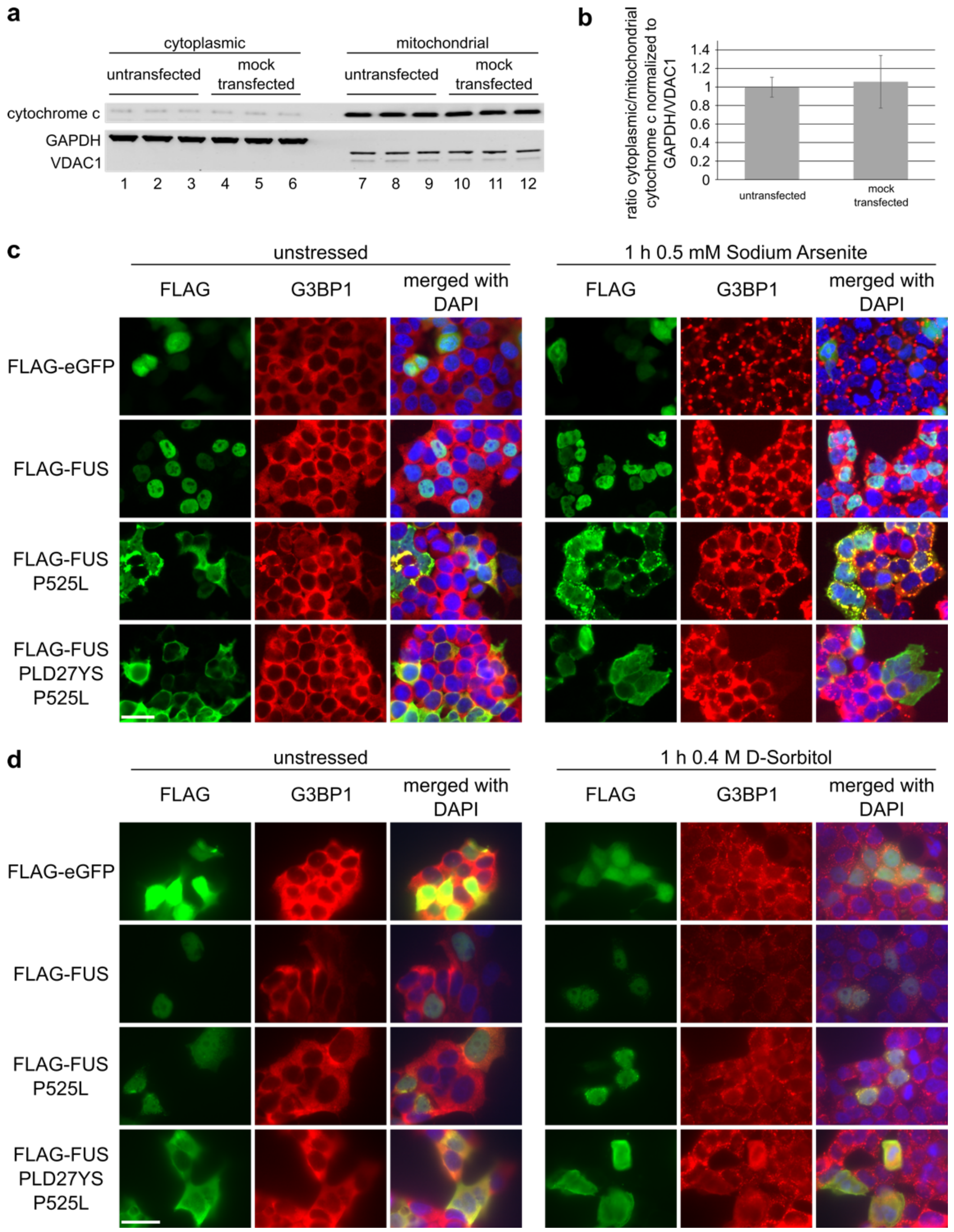


*Supplementary Figure 5 related to Figure 5*

**a** Cytoplasmic/mitochondrial fractionation of untransfected and mock transfected HEK293T cells, respectively. Cytoplasmic (lanes 1-6) and mitochondrial (lanes 7-12) fractions were analysed by western blotting using anti cytochrome c antibody (top row). GAPDH and VDAC1 (lower row) served as controls for cytoplasmic and mitochondrial fractions, respectively. **b** Quantification of cytochrome c levels in a. Shown are the ratios of cytoplasmic to mitochondrial cytochrome c relative to the first replicate of the untransfected condition. Average values and standard deviations from three biological replicates are shown. **c** Immunostaining of unstressed (left) and stressed (right) HEK293T cells transiently transfected with the indicated FLAG-FUS constructs. In addition, a FLAG-eGFP control was included. Cells were stained for FLAG (green) and the stress granule marker G3BP1 (red). Cells were counterstained using DAPI. Scale bar = 30 µm. **d** Same as in c, but with osmotic instead of oxidative stress using D-sorbitol. Scale bar = 30 µm.

| | Gene | fold change droplets vs co-IP | FDR | abundance in droplets [ppm] | abundance in co-IP [ppm] |
|---|---|---|---|---|---|
| mediator complex | MED17 | Inf | 0.2484996 | 0.390832 | 0 |
| | MED14 | Inf | 0.2289878 | 0.294638 | 0 |
| | MED12 | Inf | 0.08755816 | 1.94505 | 0 |
| | MED24 | Inf | 0.03518069 | 0.8961252 | 0 |
| | MED23 | Inf | 0.2138961 | 3.347663 | 0 |
| | MED1 | Inf | 0.02249834 | 0.7839777 | 0 |
| RNA Polymerase I | POLR1A | 9.746799 | 0.003839289 | 19.69436 | 2.020597 |
| | POLR1B | 0.460853 | 0.05772821 | 2.961577 | 6.426294 |
| | POLR1C | 7.097573 | 0.000389579 | 115.1318 | 16.22129 |
| | POLR1E | Inf | 0.05967345 | 6.569027 | 0 |
| RNA Polymerase II | POLR2A | 18.07852 | 0.006004162 | 28.18542 | 1.559056 |
| | POLR2B | 6.908363 | 0.03257622 | 30.4071 | 4.401492 |
| | POLR2C | Inf | 0.003309718 | 24.0212 | 0 |
| | POLR2E | Inf | 0.01547834 | 54.65555 | 0 |
| | POLR2H | Inf | 0.03534031 | 16.97889 | 0 |
| | POLR2I | 0 | 0.03399944 | 0 | 21.13068 |
| RNA Polymerase III | POLR3A | 240.4713 | 0.02766891 | 12.2363 | 0.05088465 |
| | POLR3B | 43.79838 | 0.08332625 | 6.157186 | 0.1405802 |
| | POLR3C | Inf | 0.009059487 | 5.164592 | 0 |
| | POLR3E | Inf | 0.4954505 | 0.5857161 | 0 |

*Supplementary Figure 6 – data extracted from Supplementary Table 1*

Mass spectrometry data extracted from Supplementary Table 1. Protein components of the mediator complex and RNA Polymerase II are more abundant under LLPS (droplets) than under non-LLPS (co-IP) conditions together with FUS. Note that most of the proteins were not detected in the non-LLPS condition (abundance of 0). Interestingly, the same appears to be true for RNA Pol I and especially RNA Pol III.

a

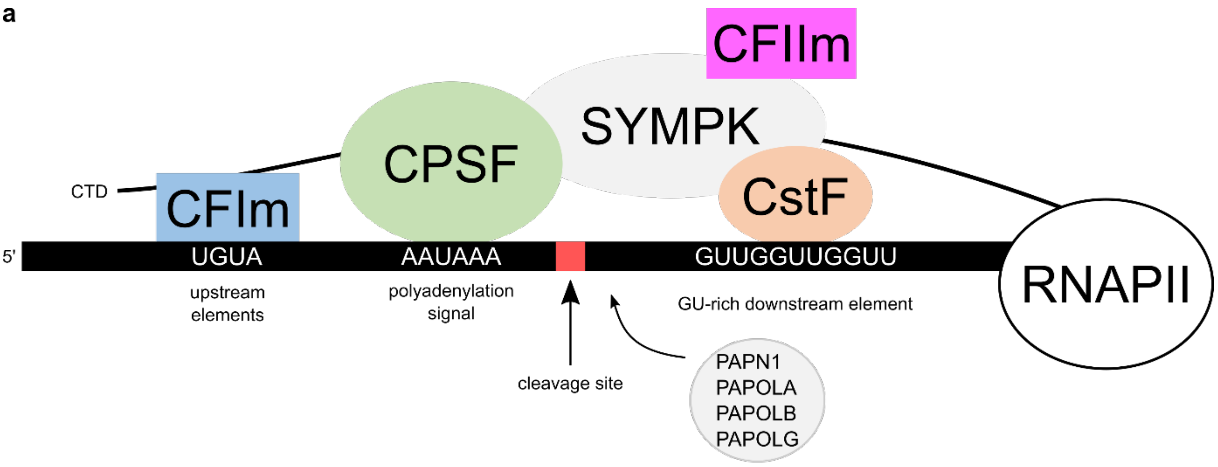


b

| | Gene | fold change droplets vs co-IP | FDR | abundance in droplets [ppm] | abundance in co-IP [ppm] |
|---|---|---|---|---|---|
| **Cleavage factor Im (CFIm)** | NUDT21 | 7.954754 | 6.54E-04 | 1810.49 | 227.5985 |
| | CPSF6 | 9.126611 | 9.64E-05 | 1136.175 | 124.4904 |
| | CPSF7 | 52.14858 | 1.77E-03 | 552.5726 | 10.59612 |
| **Cleavage and polyadenylation specificity factor (CPSF)** | CPSF1 | 0.1574387 | 2.91E-02 | 24.45202 | 155.3114 |
| | CPSF2 | 0.2187055 | 2.09E-02 | 49.26569 | 225.2604 |
| | CPSF3 | 0.3904558 | 9.57E-02 | 34.78517 | 89.08864 |
| | CPSF4 | 0 | 9.03E-02 | 0 | 27.19969 |
| | WDR33 | 0.09601287 | 1.10E-02 | 5.513332 | 57.42285 |
| | FIP1L1 | 0.256468 | 4.76E-03 | 32.53274 | 126.8491 |
| **additional factors** | SYMPK | 0.9332519 | 1.62E-01 | 44.44383 | 47.62254 |
| | PABPN1 | 0.1978951 | 2.35E-04 | 49.18344 | 248.5329 |
| | PAPOLA | Inf | 3.09E-02 | 3.763685 | 0 |
| | PAPOLB | not detected | | | |
| | PAPOLG | not detected | | | |
| **Cleavage stimulation factor (CstF)** | CSTF1 | 1.960767 | 1.16E-02 | 38.3178 | 19.54225 |
| | CSTF2 | 1.420094 | 1.74E-01 | 27.6281 | 19.45513 |
| | CSTF2T | 0.5119832 | 6.99E-01 | 0.396281 | 0.7740115 |
| | CSTF3 | 2.538518 | 1.13E-02 | 31.86525 | 12.5527 |
| **Cleavage factor IIm (CFIIm)** | PCF11 | 0.8762395 | 6.67E-01 | 1.675131 | 1.911727 |
| | CLP1 | not detected | | | |

*Supplementary Figure 7 – data extracted from Supplementary Table 1*

**a** Scheme summarizing the protein complexes involved in 3'-end processing of pre-mRNA according to (Gruber et al., 2014). **b** Mass spectrometry data extracted from Supplementary Table 1. While most components of the 3'-end processing machinery shown no clear preference for LLPS (droplets) or non-LLPS (co-IP) FUS, the three members of CFIm are strongly enriched under LLPS conditions.

### References


Gruber, A.R., Martin, G., Keller, W., and Zavolan, M. (2014). Means to an end: mechanisms of alternative polyadenylation of messenger RNA precursors. Wiley interdisciplinary reviews. RNA *5*, 183-196.

Kato, M., Han, Tina W., Xie, S., Shi, K., Du, X., Wu, Leeju C., Mirzaei, H., Goldsmith, Elizabeth J., Longgood, J., Pei, J., *et al.* (2012). Cell-free Formation of RNA Granules: Low Complexity Sequence Domains Form Dynamic Fibers within Hydrogels. Cell *149*, 753-767.

Lerner, E.A., Lerner, M.R., Janeway, C.A., Jr., and Steitz, J.A. (1981). Monoclonal antibodies to nucleic acid-containing cellular constituents: probes for molecular biology and autoimmune disease. Proceedings of the National Academy of Sciences of the United States of America *78*, 2737-2741.

Raczynska, K.D., Ruepp, M.D., Brzek, A., Reber, S., Romeo, V., Rindlisbacher, B., Heller, M., Szweykowska-Kulinska, Z., Jarmolowski, A., and Schumperli, D. (2015). FUS/TLS contributes to replication-dependent histone gene expression by interaction with U7 snRNPs and histone-specific transcription factors. Nucleic acids research *43*, 9711-9728.

Reber, S., Stettler, J., Filosa, G., Colombo, M., Jutzi, D., Lenzken, S.C., Schweingruber, C., Bruggmann, R., Bachi, A., Barabino, S.M., *et al.* (2016). Minor intron splicing is regulated by FUS and affected by ALS-associated FUS mutants. EMBO J *35*, 1504-1521.

Ruegsegger, U., Blank, D., and Keller, W. (1998). Human pre-mRNA cleavage factor Im is related to spliceosomal SR proteins and can be reconstituted in vitro from recombinant subunits. Mol Cell *1*, 243-253.

Rufener, S.C., and Muhlemann, O. (2013). eIF4E-bound mRNPs are substrates for nonsense-mediated mRNA decay in mammalian cells. Nat Struct Mol Biol *20*, 710-717.
